# Shaping sustainable harvest boundaries for marine populations despite estimation bias

**DOI:** 10.1101/2020.12.05.413070

**Authors:** Daisuke Goto, Jennifer A. Devine, Ibrahim Umar, Simon H. Fischer, José A. A. De Oliveira, Daniel Howell, Ernesto Jardim, Iago Mosqueira, Kotaro Ono

## Abstract

Biased estimates of population status are a pervasive conservation problem. This problem has plagued assessments of commercial exploitation of marine species and can threaten the sustainability of both populations and fisheries. We develop a computer-intensive approach to minimize adverse effects of persistent estimation bias in assessments by optimizing operational harvest measures (harvest control rules) with closed-loop simulation of resource–management feedback systems: management strategy evaluation. Using saithe (*Pollachius virens*), a bottom-water, apex predator in the North Sea, as a real-world case study, we illustrate the approach by first diagnosing robustness of the existing harvest control rule and then optimizing it through propagation of biases (overestimated stock abundance and underestimated fishing pressure) along with select process and observation uncertainties. Analyses showed that severe biases lead to overly optimistic catch limits and then progressively magnify the amplitude of catch fluctuation, thereby posing unacceptably high overharvest risks. Consistent performance of management strategies to conserve the resource can be achieved by developing more robust control rules. These rules explicitly account for estimation bias through a computational grid search for a set of control parameters (threshold abundance that triggers management action, *B*_trigger_, and target exploitation rate, *F*_target_) that maximize yield while keeping stock abundance above a precautionary level. When the biases become too severe, optimized control parameters– for saithe, raising *B*_trigger_ and lowering *F*_target_–would safeguard against overharvest risk (<3.5% probability of stock depletion) and provide short-term stability in catch limit (<20% year-to-year variation), thereby minimizing disruption to fishing communities. The precautionary approach to fine-tuning adaptive risk management through management strategy evaluation offers a powerful tool to better shape sustainable harvest boundaries for exploited resource populations when estimation bias persists. By explicitly accounting for emergent sources of uncertainty our proposed approach ensures effective conservation and sustainable exploitation of living marine resources even under profound uncertainty.

**Open Research Statement:** Data sets and code utilized for this research are available on Figshare. DOI: https://doi.org/10.6084/m9.figshare.13281266

## INTRODUCTION

Managers and policymakers increasingly face trade-offs in sustainably managing extractive use of living marine resources while effectively conserving biodiversity under the precautionary principle (FAO 1996, Hilborn et al. 2001, Harwood and Stokes 2003). But imperfect knowledge of social–ecological systems impedes the decision making. Scientific uncertainty (imprecision in measurements) of current population status can obscure the assessment of decline or extinction threats (Ripa and Lundberg 1996, Ovaskainen and Meerson 2010). Lack of certainty in socioeconomic dynamics that can promote noncompliance and inertia also may reduce the efficacy of management measures applied (Hilborn et al. 2001, Beddington et al. 2007, Fulton et al. 2011). If we are to achieve internationally agreed conservation targets such as sustainable use of marine resources portrayed under Sustainable Development Goal 14 (UN 2015) and Aichi Biodiversity Target 6 (CBD 2010), we must account for various sources of uncertainty (imprecision and inaccuracy) to assess overexploitation risk (Memarzadeh and Boettiger 2018) and recovery potential (Memarzadeh et al. 2019) and set conservation priorities.

In commercial capture fisheries, assessments of current population status provide a scientific basis for setting a threshold for safe harvest to prevent the decline of fish stocks. This approach may include using biological thresholds such as the population abundance that produces maximum sustainable yield (Beddington et al. 2007). The harvest of wild populations is commonly managed by applying decision rules (harvest control rules) based on such predefined thresholds to set a catch limit for the year (Beddington et al. 2007). Accurate population assessments contribute to successful implementation of management measures to sustain long-term commercial exploitation of fish populations (Hilborn et al. 2020). But systematic errors in assessments have posed a multitude of challenges (Patterson et al. 2001, Sethi 2010). If population abundance is persistently overestimated, for example, resulting overly optimistic catch advice or rebuilding plans will deplete the population, thereby threatening the sustainability of fisheries that depend on it (Walters and Maguire 1996, Memarzadeh et al. 2019). Overestimated abundance and underestimated exploitation rates, which often heighten extinction risk, have led to some historical collapses of oceanic predators (Walters and Maguire 1996, Charles 1998).

Biased estimates in perceived population status have plagued assessments of exploited marine species (Punt et al. 2020) and likely contributed to overharvest and depletion including stocks that are considered well-monitored (Brooks and Legault 2016). Inconsistency across assessments such as persistent overestimation of abundance has led to the rejection of assessments (Punt et al. 2020). Although past research has proposed solutions to estimation bias, applying these solutions remains a challenge because the bias could originate from multiple sources (Hurtado-Ferro et al. 2015, Brooks and Legault 2016, Szuwalski et al. 2017). Incomplete knowledge of the causes behind biased estimates may lead to incorrect application of the tools, inadvertently exacerbating the problems by amplifying overharvest and depletion risks (Brooks and Legault 2016, Kraak et al. 2008, Szuwalski et al. 2017). Given serious ecological and socioeconomic implications for getting it wrong, we urgently need a procedure that provides practical guidance for explicitly evaluating robustness of management strategies and designing alternatives to inform decision making to safely harvest under uncertainty (Punt et al. 2020).

We illustrate how closed-loop simulation of resource–management systems (management strategy evaluation) can help prevent estimation bias from derailing effective management of exploited marine populations. Management strategy evaluation is a flexible decision-support tool used in fisheries management (Butterworth and Punt 1999, Smith et al. 1999) and has increasingly been applied to conservation planning in marine and terrestrial systems (Milner-Gulland et al. 2001, Bunnefeld et al. 2011). This tool is designed to evaluate the performance of candidate policy instruments through forward simulations of feedback between natural resources and management systems (policy implementation and new observation) by accounting for trade-offs among management goals of stakeholders (Punt et al. 2016). Management strategy evaluation also can assess consequences of suspected sources of bias in assessments (Szuwalski et al. 2017, Hordyk et al. 2019). Here we take this approach further: we first diagnose estimation bias (robustness testing, Cooke 1999). Then, through computational optimization of harvest control rules (Walters and Hilborn 1978, Chadès et al. 2017), our proposed method searches for robust rules by explicitly accounting for bias in perceived stock status along with process (life history parameter) and observation (survey and reported catch) uncertainties. Specifically, we evaluate how robust current management procedures are to persistent estimation bias, and then demonstrate how management procedures can remain precautionary through the optimization of harvest control rules to avert mismanagement–setting overly optimistic catch limits that promote stock depletion and a future fishery closure.

## METHODS

### Management strategy evaluation framework

We simulated population and harvest dynamics, surveys, assessments, and implementation of management strategies to explore trade-offs in achieving conservation-oriented (minimizing overexploitation risk) and harvest-oriented (maximizing yield) goals through management strategy evaluation. We made use of the framework developed and adopted for commercially harvested species in the Northeast Atlantic including four North Sea demersal fish stocks (ICES 2019c) and Atlantic mackerel (*Scomber scombrus*, ICES 2020c). The framework consists of submodels that simulate 1) true population and harvest dynamics at sea (operating model, OM), from which observations through monitoring surveys and catch reporting (data generation) are made, and 2) management processes–assessments based on observations from the surveys and reported catch and subsequent decision making (management procedure, MP) (Fig. 1a, Punt et al. 2016). We used the North Sea population of saithe (*Pollachius Virens*) (ICES statistical areas: Subareas 4 and 6 and Division 3a, ICES 2019c), a demersal (bottom-water) predatory fish harvested commercially by more than a dozen European nations, as a real-world case study. And we used the State-space Assessment Model (SAM, Nielsen and Berg 2014) as estimation model (EM) and harvest control rule set for saithe (ICES 2019c); model settings and forecast assumptions are fully described in ICES (2019c). We performed all simulations in R (version 3.60, R Development Core Team 2019) using the mse R package (https://github.com/flr/mse) (ICES 2019c), part of the Fisheries Library in R (FLR, Kell et al. 2007).

**Figure 1.**
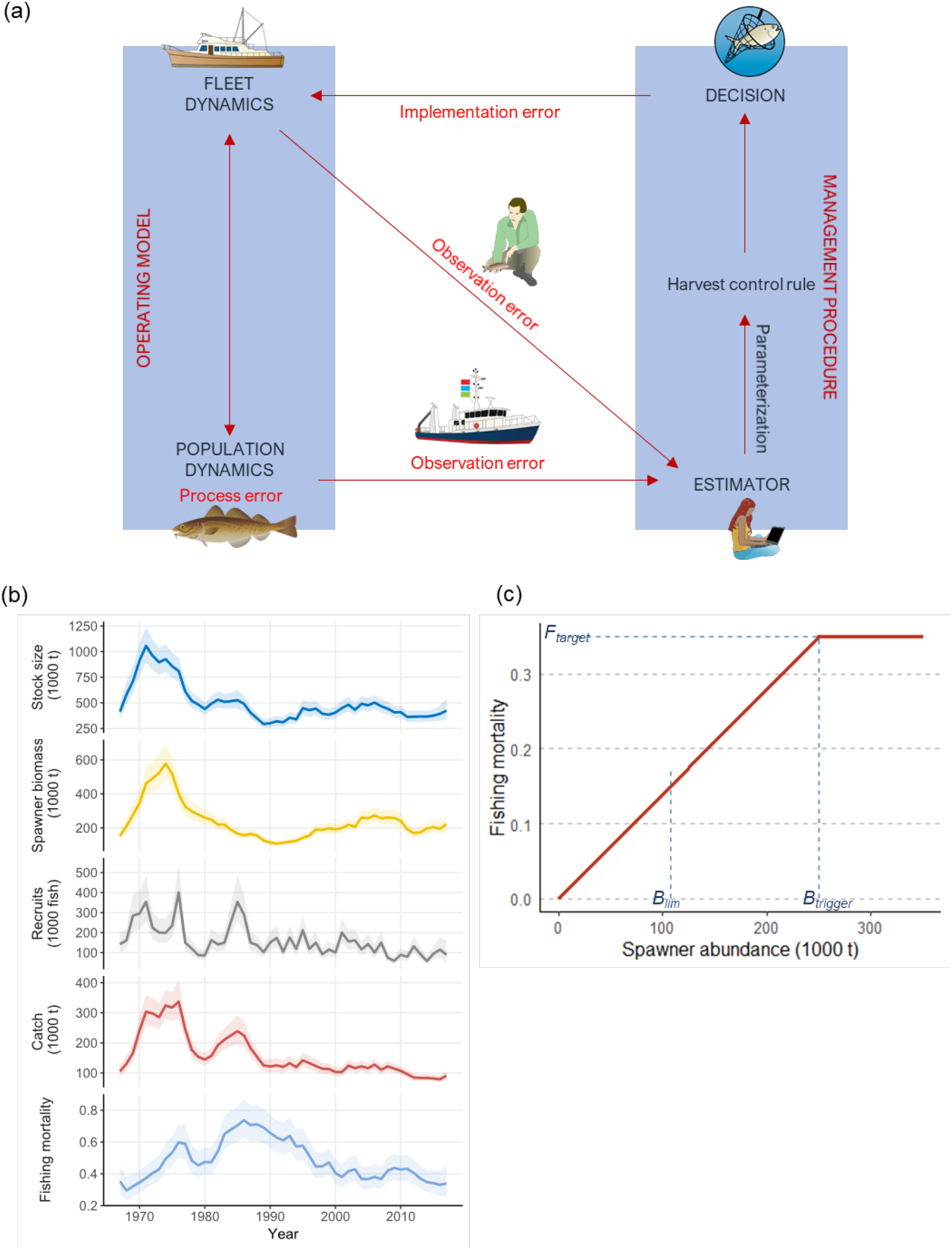
Management strategy evaluation framework and historical population and harvest dynamics of North Sea saithe. (a) Schematic of the management strategy evaluation framework (Fisheries Library in R/Assessment for All or FLR/a4a, redrawn from https://github.com/ejardim) adopted for evaluation of saithe management strategies. (b) Reconstructed saithe population and harvest dynamics taken from the 2018 assessment (ICES 2019a). Ribbons indicate 95% confidence intervals. (c) Harvest control rule evaluated in this study. Blue dashed (horizontal and vertical) lines show the harvest control rule parameters set for saithe: *B*_trigger_ = 250,000 t and *F*_target_ = 0.35 (ICES 2019c).

### Population dynamics

To simulate future population dynamics of target species, the framework uses an age-structured population model that accounts for environmental stochasticity. For saithe we modeled the population dynamics of four-year-olds and older as

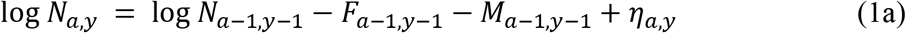

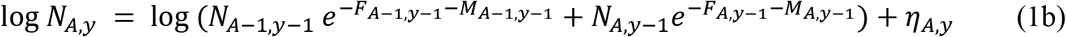

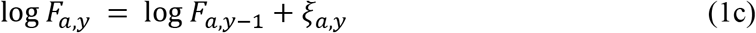

where *N_a,y_, N_a,y-1_, F_a,y_, F_a,y-1_, M_a,y_*, and *M_a,y-1_* are a-year-old numbers, fishing mortality rates, and natural mortality (non-fishing such as starvation and diseases) rates in year *y* and *y*-1, and *η_a,y_* and *a,y* are multivariate normally distributed variables, reflecting process errors correlated between ages within years (Appendix S2: Fig. S1, Nielsen and Berg 2014). *F_a,y_*-1 is time-varying and simulated through the implementation of harvest control rules (see *Management procedure* below). Historical surveys indicate that 10-year-olds and older are relatively uncommon, and we simulated them as a dynamic aggregate pool (known as a plus group in fishery stock assessment, *N_A_, F_A_*, and *M_A_*).

We simulated density-dependent regulation of recruitment in the population dynamics with a segmented regression (ICES 2019c) relating adult biomass to the number of recruits (three-year-olds for saithe) as

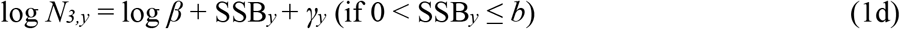

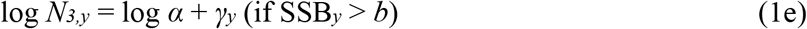

where SSB*y* is adult biomass (known as spawning stock biomass, t) in year *y*, which is the sum of the product of age-specific numbers, masses, and maturity rates, *β, b*, and *α* are parameters, and *γ_y_* is process error in year *y*.

We developed the OM using data and life history parameter estimates taken from the 2018 assessment (Fig. 1b, ICES 2018), which represents the best available information on the past (1967–2017) population and harvest dynamics (Fig 1b and Appendix S1). The data sources, survey methods, and model structure have been extensively documented in ICES (2016) and ICES (2019a). Briefly, we parameterized the model with 51-year estimates of age-specific masses (g, Appendix S1: Table S3–S4) and maturity rates (proportion of adults, Appendix S1: Table S5), and natural mortality rates assumed at 0.2 year^−1^ for all ages and years. Then, we fitted the population model to time series data of commercial catch (age-aggregated biomass of German, French, and Norwegian trawlers in 2000–2017, tonnes or t, Appendix S1: Table S6 and Appendix S2: Fig. S1) and age-specific (ages three to eight) abundance indices (International bottom trawl surveys in the third quarter, IBTS-Q3, in 1992–2017, Appendix S1: Table S7 and Appendix S2: Fig S2) (ICES 2018) using SAM (see *Monitoring and catch surveys* below for details of computing catch and age-specific abundance indices).

We projected true population and catch dynamics annually for 21 years (2018–2038). To account for process uncertainty (year-to-year variability in survival rate), we generated 1000 realizations of stochastic populations using the variance-covariance (inverse hessian) matrix of age-specific numbers and fishing mortality rates taken from the 2018 assessment (Appendix S2: Fig. S3a, ICES 2019c). We derived a set of mean age-specific masses, maturity rates, and fishing gear selectivity by randomly selecting a year with replacement from the 2008–2017 data; this process was repeated independently for each replicate every year to account for environmental stochasticity.

To account for environmental stochasticity in density-dependency of recruitment, we first parameterized the spawner–recruit model by fitting it to the 1998–2017 data on SSB and recruit numbers by resampling residuals with replacement. Because preliminary analyses had revealed gaps in the resampling process (ICES 2019c), we used a kernel density function to smooth the resulting distribution of residuals from the fitted regression. Then, we resampled residuals from the distribution and applied these to model outputs to generate recruits every year (Appendix S2: Fig. S4a,b); this process was repeated independently for each replicate. Preliminary analyses showed little evidence of temporal autocorrelation in recruitment (Appendix S2: Fig. S4c).

### Monitoring and catch surveys

We simulated future annual monitoring of the population and harvest, which are subject to error, by adding observation error to age-specific survey indices and aggregated catch computed from the OM. To simulate deviances to the observed survey index (IBTS-Q3) we used the variance-covariance matrix for the survey index to account for observation error correlated between ages (Appendix S2: Fig. S5a and S6a). Survey observations (*I*) are generated as:

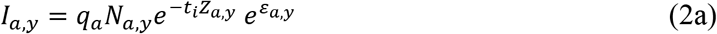

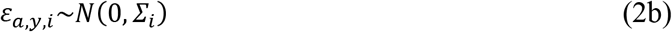

where *Z_a,y_* is *a*-year-old total (*F_a,y_* + *M_a,y_*) mortality rate in year *y* from the OM; *q_a_* are *a*-year-old survey catchabilities for the survey *i*; *t* is the timing of the annual survey (0.575 for IBTS-Q3). *ε_a,y_* represents multivariate normally distributed errors with mean zero and standard deviation *Σ* defined by the variance-covariance matrix between ages within years (ICES 2019b). Observation error is applied to age-specific abundance indices as multiplicative lognormal error (Appendix S2: Fig. S5a).

To avoid using the age information twice (once in computing age-specific catches and again in selectivities), we computed a fishable biomass index, a combined (German, French, and Norwegian trawlers) index from the OM (Appendix S2: Fig. S5b and S6b) standardized by average fishing mortality rates as:

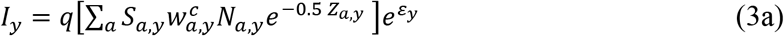

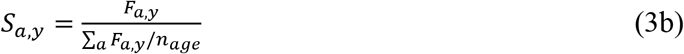

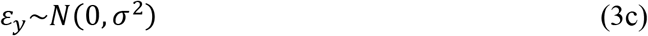

where *q* is the catchability; 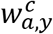 are *a*-year-old catch masses in year *y*; 0.5 indicates projection to mid-year; *S_a,y_* is the selectivity of *a*-year-olds in year *y*; *n*_age_ is the number of age classes in the population; and *ε_y_* is a normally distributed error with mean zero and standard deviation *σ* in year *y* (Appendix S2: Fig. S3c). We used a version of SAM (Nielsen and Berg 2014) accounting for this change (https://github.com/fishfollower/SAM/tree/biomassindex).

### Management procedure

The MP simulates decision making by managers based on perceived current stock status and model-based harvest control rules (Fig. 1a). The current status is assessed annually by fitting the EM to the time series (past plus most recent year, *y*) data simulated from the observation model (survey and catch data, *I_a,y_* and *I_y_*) before the provision of catch advice (May of the following year, y+1, for saithe). Under the control rule set for saithe (ICES 2019c), when the estimated SSB at the start of the advice year following the assessment year (terminal year) remains above a fixed threshold (*B*_trigger_) (Fig. 1b), the catch limit is computed based on target exploitation rate (*F*_target_). These two control parameters (*B*_trigger_ and *F*_target_) are designed to prevent overharvesting by accounting for uncertainty in population and harvest dynamics (Rindorf et al. 2016). For consistency we used the same parameter values of the control rule that had been estimated in ICES (2019c) (*B*_trigger_ = 250,000 t and *F*_target_ = 0.35, see *Population and management measure performance* below for detail). When the SSB falls below *B*_trigger_, exploitation rate is adjusted to *F*_target_ scaled to the proportion of SSB relative to *B*_trigger_ (Fig. 1c), thereby allowing the population to rebuild (adaptive harvesting). In simulations the advice year’s SSB (SSB*y*+1) is first forecasted with the EM (SAM) using the average of estimated fishing mortality rates in the most recent three years (known as *F* status quo). Then the target exploitation rate for the advice year (*F*_*y*+1_) is determined to compute the catch limit (*C*_*y*+1_) as

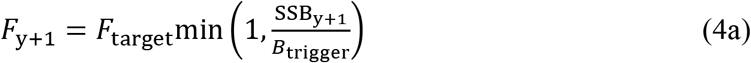

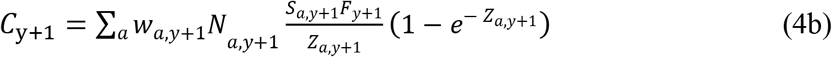

where *W_a,y+1_, N_a,y+1_, S_a,y+1_*, and *Z_a,y+1_* are as above and forecasted for the advice year.

### Population and management measure performance

We computed conservation-oriented (risk of stock depletion) and harvest-oriented (median catch and interannual catch variability, ICV) metrics averaged across 1000 replicates of short-term (2019–2023) and long-term (2029–2038) projections from the OM to evaluate performance of the harvest control rules applied. We chose the number of replicates based on the stability of risk (ICES 2019c). Risk of stock depletion is defined as the maximum annual probability of SSB falling below a limit threshold, *B*_lim_ (Fig. 1c), a spawner abundance below which reproductive capacity of the populatio is expected to decline (Rindorf et al. 2016), consistent with previous analyses (ICES 2019b). We computed the risk based on the proportion of 1000 replicates with annual estimates of SSB < *B*_lim_. The International Council for the Exploration of the Sea (ICES) defines reference points following its guidelines (ICES 2021). *B*_lim_ is set to 107,297 t for saithe (2019a) and based on the lowest observed historical SSB. Following ICES (2021), *B*_lim_ is used as the basis for computing maximum sustainable yield (MSY) *B*_trigger_ (ICES 2020a, 2021) as

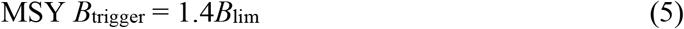

which is a default value of *B*_trigger_. *F*_MSY_ (used as default *F*_target_) is estimated with the eqsim R package (https://github.com/ices-tools-prod/msy). EqSim produces a long-term stochastic projection (ICES 2015, 2017, 2020a). The resulting control parameters follow the MSY approach but are constrained under the precautionary criteria (ICES 2021). As part of the latest management strategy evaluation both *B*_trigger_ and *F*_target_ were optimized through a grid search by maximizing median catch limits while maintaining long-term risk ≤ 0.05 (Appendix S2: Fig. S7 and S8, ICES 2019b). We computed ICV (a proportional change in catch limit) as

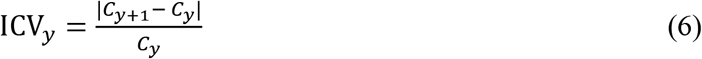

where *C_y+1_* and *C_y_* are projected catches (eq. 4b) in year *y*+1 and *y*.

### Estimation bias scenarios

To evaluate how managing with persistently biased assessments degrades performance of harvest control rules and potential to achieve management goals, we simulated hypothetical scenarios of bias in perceived spawner abundance and fishing mortality rate in annual assessments. Although bias can emerge in both directions (over- and under-estimation), they have asymmetric implications for conservation and harvest decision making by managers (Hordyk et al. 2019). We analyzed scenarios that can cause severe conservation issues for exploited species: SSB overestimation and mean *F* (averaged across four to seven-year-olds for saithe) underestimation simultaneously. We simulated six scenarios by introducing a bias (0%/baseline, 10%, 20%, 30, 40%, and 50% per year) in estimating age-specific numbers and fishing mortality rates in the terminal year of annual assessment before forecasting SSB and mean *F* and projecting a catch limit. The magnitudes of realized biases in these parameters however varied among simulations because of process uncertainty. We introduced a bias as

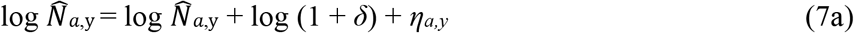

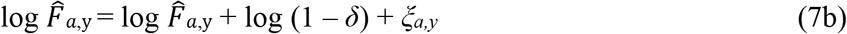

where 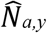 and 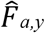 are estimated *a*-year-old numbers and fishing mortality rates in year *y* from the EM, and *δ* is a bias (in proportion). The biased estimates are then used to compute SSB_*y*+1_ prior to projecting a catch limit using the harvest control rule as above (eqs. 4a,b). Note that for simplicity and generality these bias scenarios are designed to illustrate our proposed approach to generic estimation bias in assessments, rather than specific scenarios of persistent, time-varying bias that may cumulatively emerge between assessments as input data are updated owing to model misspecification and biased input data (known as retrospective pattern, ICES 2020b, Punt et al. 2020). We analyzed all scenarios based on the performance metrics (risk, median catch, and ICV) of short-term and long-term projections.

### Developing robust management measures

To evaluate how precautionary the harvest control rule needs to be to minimize adverse effects of biased estimates in the assessment on catch advice provisioning, we explored alternative values of the two control parameters of the harvest control rule (*B*_trigger_ and *F*_target_) and projected catch limits under the same bias scenarios (overestimated SSB and underestimated mean *F*) through management strategy evaluation. Building on the grid search from the latest evaluation (ICES 2019c) and using *B*_trigger_ = 250,000 t and *F*_target_ = 0.35 as baselines, we explored a finite number of select candidate combinations of the parameters (12 *B*_trigger_ × 16 *F*_target_ = 192 per scenario or 1,920,000 unique runs in total) for reoptimization to illustrate our proposed approach. We conducted a restricted grid search in parameter spaces of *B*_trigger_ (210,000 to 320,000 t with 10,000 t increments) and *F*_target_ (0.24 to 0.39 with 0.01 increments) for each bias scenario. We computed median catch limits and risk from the simulations and optimized the parameter sets by maximizing median catch limits while maintaining long-term risk ≤ 0.05.

## RESULTS

### Performance of harvest measures with estimation bias

An increasing amount of estimation bias in annual assessments was found to increase median catch and overharvest risk in the short term. Although median SSBs declined by as much as 30% in the OM (Fig. 2a), with SSB overestimation, median catches rose by 15–44% relative to the baseline (Fig. 3a), increasing mean *F*s in the OM by 19–80%, which were underestimated in the EM by on average 42% (Fig. 2b). As a result, biased assessments elevated risks as much as 17-fold (Fig. 3a). Mean ICV responded nonlinearly to biased estimates, and the distribution was highly skewed (Fig. 3a).

**Figure 2.**
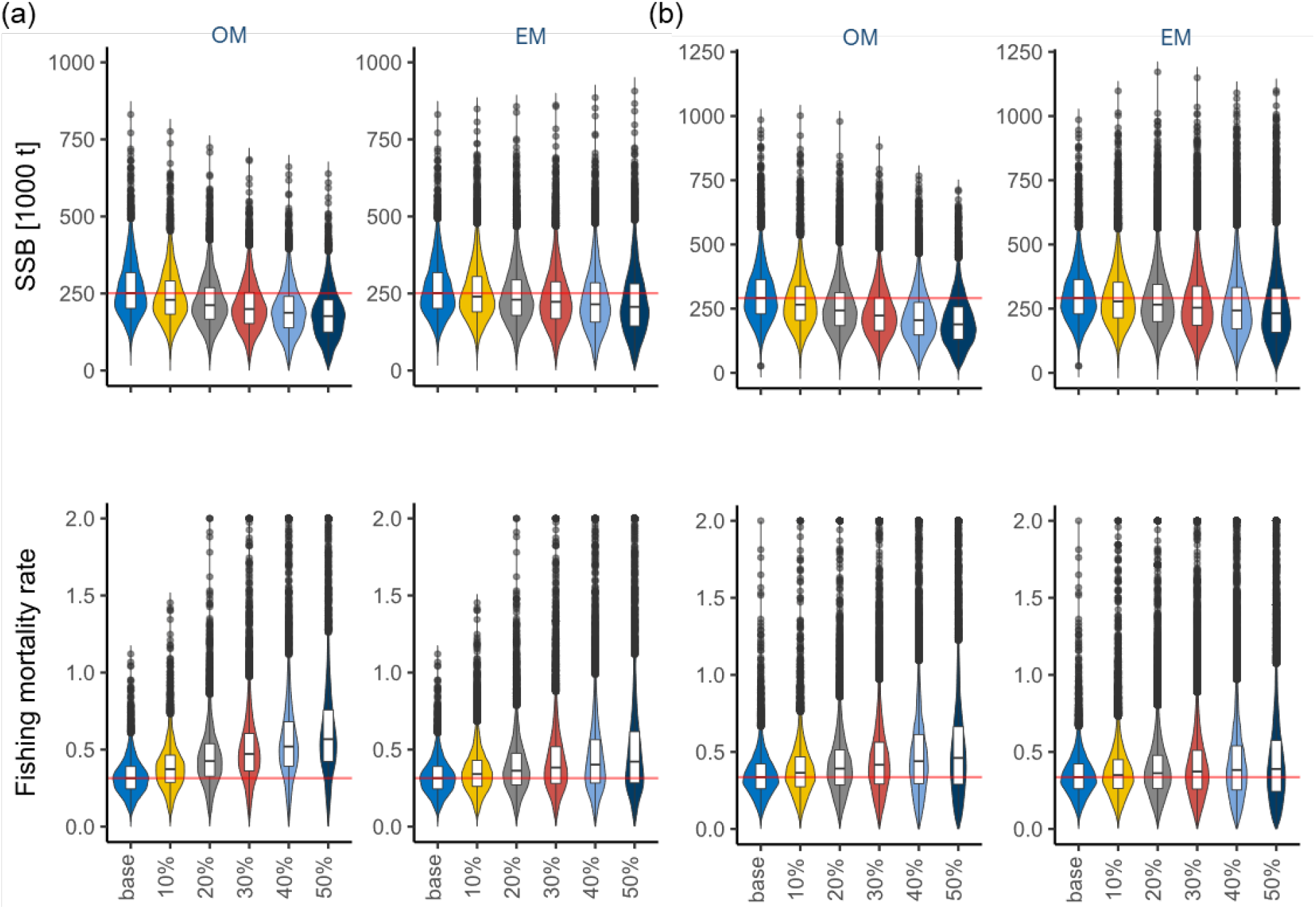
Stock abundance (SSB) and fishing pressure of North Sea saithe from the population operating and estimation models (OM and EM) under scenarios of varying levels of estimation bias: (a) short-term (2018–2023) and (b) long-term (years 2029–2038). Violin plots indicate frequency distributions of performance metrics. Horizontal lines (from bottom to top) within the box plots indicate the 25th, 50th, and 75th percentiles; whiskers (of the box plots) extend to the largest and smallest values within 1.5x the inter-quartile range (IQR) from the box edges; and black circles indicate the outliers. Fishing mortality rates are computed by averaging across age-specific fishing mortality rates of four to seven-year-olds. Red horizontal lines indicate median values from the baseline scenario.

**Figure 3.**
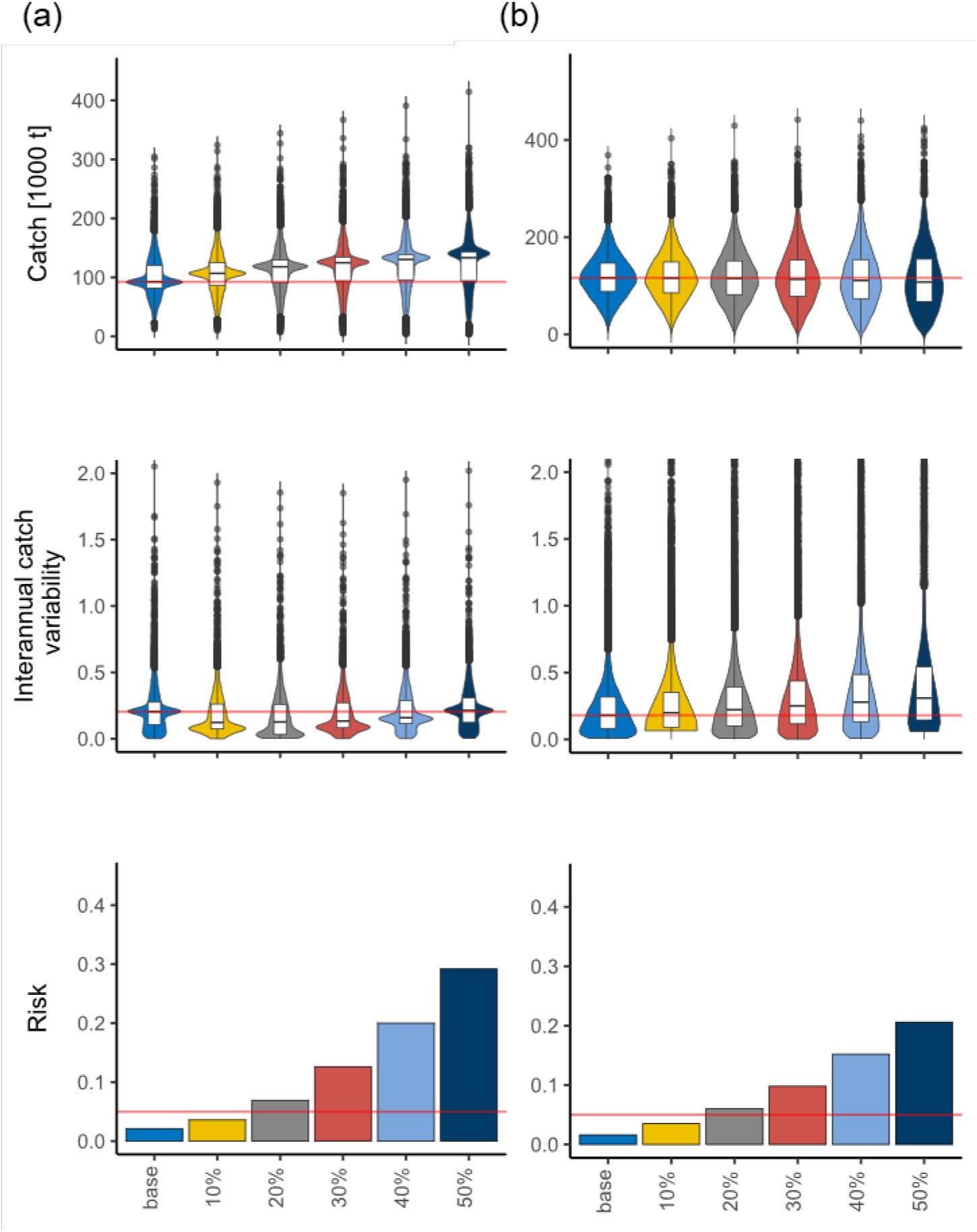
Performance of the harvest control rule for North Sea saithe under six scenarios of varying levels of estimation bias (overestimation of stock abundance and underestimation of fishing mortality rate): (a) short-term (2018–2023) and (b) long-term (years 2029–2038). The performance was evaluated with median catch (t), interannual catch variability (ICV), and risk. Risk is the maximum probability of SSB falling below *B*_lim_ (107,297 t). Violin plots indicate frequency distributions of performance metrics. Horizontal lines (from bottom to top) within the box plots indicate the 25th, 50th, and 75th percentiles; whiskers (of the box plots) extend to the largest and smallest values within 1.5x the inter-quartile range (IQR) from the box edges; and black circles indicate the outliers. Red horizontal lines indicate median values from the baseline scenario (catch and ICV) or the precautionary threshold (risk = 0.05).

In the long-term the estimation bias was found to increase ICV and risk but had negligible effect on median catch. Biased estimates reduced median SSB in the OM by as much as 35% (resulting in a 37% increase in mean *F*) relative to the baseline; this reduction was underestimated in the EM by on average 53% (Fig. 2a,b). With overestimated SSBs and largely unadjusted *Ftarget*, median catches remained unchanged (~113,000 t, Fig. 3b). Also, biased assessments amplified temporal variations (CVs in medians of replicates) in both SSB and mean *F* in the OM as much as ~71%, thereby increasing ICVs by up to 72%, which, combined with reduced SSBs, elevated risks 2–13-fold (Fig. 3b).

### Harvest control rule optimization

The proportion of the select grid search area evaluated through management strategy evaluation that remained precautionary (which we define as safe harvest margin) progressively shrank as more bias was introduced (Fig. 4 and Table 1). Within the safe harvest margin, the fishery yielded highest catches at lower (by 0.02–0.10) *F*_target_ and higher (by 10,000–60,000 t) *B*_trigger_ (Table 1 and Fig. 4). With reoptimization of these control parameters the control rule was projected to produce higher (by 6.7–25%) short-term catches and maintain similar (<3.0% deviation from the baseline) long-term catches under all bias scenarios (Table 1). And both short- and long-term SSBs declined by 3.1–6.9% and long-term ICVs rose by less than 1.5% (Table 1).

**Figure 4.**
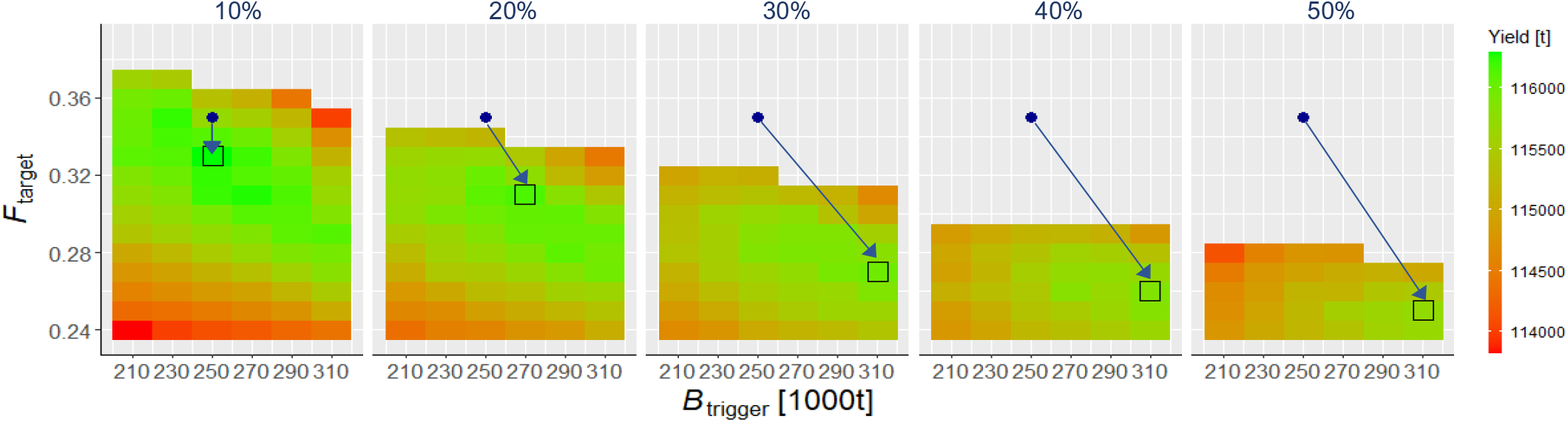
Grid search for combinations of the harvest control rule parameters (*F*_target_ and *B*trigger) for North Sea saithe under five scenarios of varying levels of estimation bias (overestimation of stock abundance and underestimation of fishing mortality rate). Heat maps indicate median catch for only combinations that meet the precautionary criterion (risk ≤ 5%) in the long term (years 2029–2038). Black rectangles indicate combinations of the harvest control rule parameters with the highest median catch. Blue circles indicate the parameter sets optimized without estimation bias (*B*_trigger_ = 250,000 t and *F*_target_ = 0.35, ICES 2019c).

**Table 1.** Optimized control parameters (*F*_target_ and *B*_trigger_)^†^ of the harvest control rule set for North Sea saithe and performance metrics^‡^ from management strategy evaluation under scenarios of varying levels of estimation bias in assessments.

| scenario<sup>§</sup> | $F_{\text{target}}$ | $B_{\text{trigger}}$ | short-term (2019–2023) | | | | long-term (2029–2038) | | | | |
| --- | --- | --- | --- | --- | --- | --- | --- | --- | --- | --- | --- |
|  |  |  | Catch | ICV | SSB | risk<sup>¶</sup> | catch | ICV | SSB | risk<sup>¶</sup> | SHM<sup>¶</sup> |
| base | 0.35 | 250000 | 92464 | 20 | 251973 | 2.0 | 116700 | 17.7 | 292067 | 1.5 | - |
| 10% | 0.33 | 250000 | 101786 | 13 | 238194 | 3.2 | 116288 | 17.8 | 279135 | 2.5 | 84.4 |
| 20% | 0.31 | 270000 | 103545 | 13 | 235356 | 3.3 | 116154 | 18.7 | 274958 | 3.0 | 65.6 |
| 30% | 0.27 | 310000 | 93047 | 20 | 252123 | 2.2 | 115984 | 18.0 | 293711 | 2.2 | 53.1 |
| 40% | 0.26 | 310000 | 101131 | 14 | 240643 | 2.9 | 115863 | 18.4 | 282929 | 2.5 | 37.5 |
| 50% | 0.25 | 310000 | 104943 | 12 | 234525 | 3.3 | 115730 | 19.1 | 274228 | 2.8 | 29.2 |

<sup>†</sup>The model parameters were optimized for the highest median catch while meeting the precautionary criterion: long-term risk $\leq 5\%$ (ICES 2019c).
<sup>‡</sup>The performance was evaluated with short-term and long-term median catch (t), interannual catch variability (%), ICV, median spawning stock biomass (SSB, t), and risk (%).
<sup>§</sup>Scenarios simulate SSB overestimation and mean (averaged across four to seven-year-olds) fishing mortality rate ( $F$ ) underestimation.
<sup>¶</sup>Risk is the maximum probability of SSB falling below $B_{\text{lim}}$ (107,297 t) over a given period. Safe harvest margin (SHM) indicates the proportion (%) of the grid-search area with the harvest rules that remain precautionary (Fig. 4).

## DISCUSSION

An optimization approach applied through management strategy evaluation offers a powerful decision-support tool to develop robust harvest control rules for sustainable fisheries even when severe estimation bias persists in assessments. For North Sea saithe, increasingly severe biases (abundance overestimation and fishing pressure underestimation) initially set overly optimistic catch limits that deplete the stock. But unacceptably high long-term risks of missing management targets result from progressively amplified fluctuations in annual catch limits. With computational optimization our proposed approach can help develop harvest control rules to achieve robust, cost-effective performance: low risks and stable catch limits–less disruption to fishing communities. By explicitly accounting for persistent estimation bias in assessments this approach can guide resource managers in balancing the trade-off in managing commercial exploitation: achieving stability in harvest while also maintaining sustainable resource populations.

### Costs of managing with estimation bias

How robust management measures are to biased estimates in assessments would depend on life history, fishing operation, and current status of a given species or population (Hurtado-Ferro et al. 2015, Wiedenmann and Jensen 2018, Hordyk et al. 2019). Our North Sea saithe case study is based on the 2018 assessment in which the stock is in good condition (~37% above MSY *B*_trigger_, ICES 2019c). Analyses show the current harvest control rule is robust to a moderate amount of bias (up to ~16%, based on our further analyses with 1% increments between 10% and 20%) in assessments and the stock can be sustainably managed at an acceptable level of risk (≤5% probability of stock depletion). Simulations revealed, however, that managing harvest with more severely biased assessments can progressively amplify the risk of overharvesting but the causes of heightened risk vary over time. The risk initially increases as the population becomes depleted owing primarily to overly optimistic projections of annual catch limits. Past research suggests that this pattern can emerge from misspecification of an estimation model such as unaccounted temporal variability in demographic parameters (Szuwalski et al. 2017) and overestimated natural mortality rate (Hordyk et al. 2019), and biased input data such as underreported catch (Hordyk et al. 2019). Our exploratory analyses with misspecified natural mortality rates also show that assessments with an overestimated (by 50%) natural mortality rate can underestimate fishing pressure and overestimate stock size, increasing the risk of depletion (by 67%, Appendix S2: Fig. S9). Over time managing with biased assessments would destabilize the stock, which is displayed as amplified variations in both stock abundance and fishing pressure in our case study. Yields also would become increasingly more variable (by as much as 74% for saithe), elevating the probability of overharvesting. Even when the long-term risk of managing with estimation bias remains within acceptable levels (under <20% bias scenarios in our case study), harvesting destabilized stocks may have more uncertain consequences for population persistence and yield.

Large year-to-year fluctuations in catch limit are disfavored by fishing communities (Anderies 2015) and a management measure to suppress the fluctuations (known as stability or catch constraint) is commonly applied in industrial exploitation (ICES 2019b). But evidence for the efficacy of this policy tool remains limited (but see Kell et al. 2005, Kell et al. 2006, Goto et al. 2021) especially when assessments suggest persistent biases in stock status. Applying the fluctuation-suppressing measure may, to some extent, limit catch variability inflated by managing with biased assessments. But the risk of stock depletion likely remains unacceptably high because this tool may not be sufficiently sensitive to rapid population declines and unlikely prompts large enough reductions in annual catch limit effectively (Kell et al. 2005, Kell et al. 2006, Goto et al. 2021).

The time-varying consequences of biased estimates in assessments also may present a dilemma for managers in decision making, as illustrated for several exploited marine species (Deroba 2014, Hordyk et al. 2019). Managing with biased assessments would produce higher yields (and revenues) in the short term but would amplify catch fluctuations and thus probabilities of depletion in the long term. Trade-offs between short-term gains and long-term losses (or vice versa) are common dilemmas in managing natural resources (Mangel et al. 1996, Carpenter et al. 2015). Past research focuses on developing solutions to biased assessments in fisheries management (Brooks and Legault 2016, Wiedenmann and Jensen 2018). Capturing how managers and fishing communities respond to these changes also would contribute to developing effective strategies for sustainable use of resource populations (Fulton et al. 2011). For example, historical records tell us that realized catch limits and landings in the Northeast Atlantic on average varied less than recommended by scientific advice (Patterson and Résimont 2007), which may attenuate or amplify the effects of biased assessments on the sustainability of harvesting. In situations where the science that management advice is based on becomes increasingly unreliable, evaluating both short- and long-term consequences of taking certain management actions would aid managers make decisions effectively. Our findings reemphasize alternative harvest measures need to be explicitly assessed before implementation when giving a scientific basis to inform defensible decision making.

### Managing risks under rising uncertainty

Our analyses suggest persistent overestimation of abundance and underestimation of fishing pressure can mask the extent of overharvesting and depletion, thereby delaying management responses (asynchronized resource–fishery dynamics, Fryxell et al. 2010). Although a certain time lag in the management cycle (from monitoring surveys to provisioning of catch advice) is unavoidable, severe estimation bias can promote management inertia. Once population abundance reaches a biological limit threshold (*B*_lim_ for example), the population may even become unresponsive to any measure for recovery (Allee effect, Kuparinen et al. 2014). One proposal to minimize adverse effects of estimation bias is by identifying the sources of and correcting for model misspecification such as accounting for time-varying demographic parameters in an estimation model (Szuwalski et al. 2017). But without prior knowledge of true demographic processes of the population the current form of this method may not sufficiently reduce bias or may even exacerbate the problem if incorrectly applied (Szuwalski et al. 2017). Also, if biases originate from two or more demographic parameters, uncertainties in these misspecified parameters may covary and interact unpredictably, making the application of the method challenging for many harvested populations.

To circumvent this challenge others suggest annual catch limits be proportionally adjusted using an index that quantifies relative deviation in population metrics (such as stock abundance) between assessments (known as Mohn’s *ρ*) (Deroba 2014, Brooks and Legault 2016). Although this index can be useful as a diagnostic, past analyses suggest the index may not necessarily reflect the magnitude and direction of bias (Hurtado-Ferro et al. 2015, Brooks and Legault 2016, Wiedenmann and Jensen 2018). When applied the outcomes and net benefits can be equivocal in both the short- and long-terms (Deroba 2014, Brooks and Legault 2016).

Shifting the focus from assessment to decision making in management strategy evaluation (Fig. 1a), our analysis shows the undesirable outcomes of managing with biased assessments can be avoided by developing more precautionary measures to set annual catch limits through dynamic optimization of the control parameters of harvest control rules. For our saithe case, when estimation bias becomes too severe, lowering target exploitation rate and raising threshold abundance that trigger management action–early intervention–would maintain not only low probabilities of stock depletion (<3.5% when SSB < *B*_lim_) (and thus a fishery closure) but also short-term catch stability (<20% year-to-year variation) without foregoing yields, thereby minimizing disruption to fishing communities. Although this approach needs to be tested with more case studies, our work demonstrates the optimization approach can guide managers in making decisions to cost-effectively safeguard against ecologically and socioeconomically undesirable outcomes of managing risks with biased assessments.

Like all model-based methods our proposed approach also has limitations. The main aim of this work was to develop an alternative approach to guide resource managers in decision making to support sustainable use of resource populations despite estimation bias. For this reason, we did not explore underlying mechanisms of the bias propagating through a resource–management system. Analyses show that even with optimization our ability to safely harvest the populations would become progressively limited (less margin of error in setting the precautionary harvest rules or “safe operating space”, Carpenter et al. 2015) as the magnitude of bias increases. We encourage continued efforts to develop methods to identify root causes of bias and to minimize their adverse effects on scientific advice (Hurtado-Ferro et al. 2015, Szuwalski et al. 2017, Hordyk et al. 2019).

Another caveat of our approach is computational intensity (requiring extensive parallel computing on a high-performance computer cluster), which may pose challenges in its application especially for more complex management objectives (more control parameters) (Walters and Hilborn 1978, Chadès et al. 2017). Methods have been recently adopted to improve the efficiency of computational optimization including genetic algorithms (Fischer et al. 2021), partially observable Markov decision process (Memarzadeh & Boettiger 2018), stochastic process (Wiedenmann et al. 2015), bootstrapping (ICES 2020a), and Bayesian statistics (ICES 2020a). Future research would benefit from applying these techniques to expand this feedback-based approach to tackling estimation bias in assessment.

More broadly, our proposed approach using management strategy evaluation, which is designed to account for multiple sources of uncertainty (Punt et al. 2016), offers a robust alternative to managing resource populations when biases in assessments persist. This approach can not only act as a diagnostic to evaluate the robustness of management measures by explicitly accounting for long-term (a decade or more) consequences but also present an adaptive, transparent way to improve protective measures when the perception deviates too far from reality. Given ubiquity of estimation bias and challenges in identifying the sources (Hurtado-Ferro et al. 2015, Brooks and Legault 2016, Szuwalski et al. 2017) we suggest the bias be routinely evaluated through management strategy evaluation as an additional source of uncertainty, and harvest control rules be (re)optimized when the bias becomes too severe.

Demand for wild-capture fisheries, which provide food, nutrition, and job security, will continue to rise with growing human populations in the coming decades (Costello et al. 2020). Changing ocean conditions are also projected to increase environmental stochasticity, amplifying resource population and harvest fluctuations (Brooks and Legault 2016). Higher environmental stochasticity may promote autocorrelation in population fluctuation (Ripa and Lundberg 1996, Gamelon et al. 2019) and amplify the magnitude of assessment error, thereby further shrinking safe harvest margins. These anticipated issues underscore greater needs for taking precautionary measures in shaping resilient management policies (adopting “resilience-thinking”, Fischer et al. 2009) to safeguard shared resources in the face of rising uncertainty.

## Acknowledgements

We thank all participants of the ICES Workshop of North Sea Management Strategies Evaluation (WKNSMSE) for feedback on the saithe management strategy evaluation work. We especially thank Anders Nielsen for assistance on SAM. We also thank Chris Legault and anonymous reviewers for comments on earlier versions of the manuscript. Some figures use images from the IAN Symbols, courtesy of the Integration and Application Network, University of Maryland Center for Environmental Science (ian.umces.edu/symbols/). This project was partially funded by the Institute of Marine Research’s REDUS (Reduced Uncertainty in Stock Assessments) project.

## Literature Cited

Anderies, J. M. 2015. Managing variance: key policy challenges for the Anthropocene.Proceedings of the National Academy of Sciences 112:14402–14403.

Beddington, J. R., D. J. Agnew, and C. W. Clark. 2007. Current problems in the management of marine fisheries. Science 316:1713–1716.

Brooks, E. N., and C. M. Legault. 2016. Retrospective forecasting—evaluating performance of stock projections for New England groundfish stocks. Canadian Journal of Fisheries and Aquatic Sciences 73:935–950.

Bunnefeld, N., E. Hoshino, and E. J. Milner-Gulland. 2011. Management strategy evaluation: a powerful tool for conservation? Trends in Ecology & Evolution 26:441–447.

Butterworth, D., and A. Punt. 1999. Experiences in the evaluation and implementation of management procedures. ICES Journal of Marine Science 56:985–998.

Carpenter, S. R., W. A. Brock, C. Folke, E. H. Van Nes, and M. Scheffer. 2015. Allowing variance may enlarge the safe operating space for exploited ecosystems. Proceedings of the National Academy of Sciences 112:14384–14389.

CBD. 2010. The strategic plan for biodiversity 2011-2020 and the Aichi Biodiversity Targets. COP 10 Decision X/2. CBD, Montreal, Canada.

Chadès, I., S. Nicol, T. M. Rout, M. Péron, Y. Dujardin, J.-B. Pichancourt, A. Hastings, and C.E. Hauser. 2017. Optimization methods to solve adaptive management problems. Theoretical Ecology 10:1–20.

Charles, A. T. 1998. Living with uncertainty in fisheries: analytical methods, management priorities and the Canadian groundfishery experience. Fisheries Research 37:37–50.

Cooke, J. 1999. Improvement of fishery-management advice through simulation testing of harvest algorithms. ICES Journal of Marine Science 56:797–810.

Deroba, J. J. 2014. Evaluating the Consequences of Adjusting Fish Stock Assessment Estimates of Biomass for Retrospective Patterns using Mohn’s Rho. North American Journal of Fisheries Management 34:380–390.

FAO. 1996. Precautionary approach to capture fisheries and species introductions. FAO Technical Guidelines for Responsible Fisheries 2, FAO.

Fischer, J., G. D. Peterson, T. A. Gardner, L. J. Gordon, I. Fazey, T. Elmqvist, A. Felton, C. Folke, and S. Dovers. 2009. Integrating resilience thinking and optimisation for conservation. Trends in Ecology & Evolution 24:549–554.

Fischer, S. H., J. A. A. De Oliveira, J. D. Mumford, and L. T. Kell. 2021. Using a genetic algorithm to optimize a data-limited catch rule. ICES Journal of Marine Science 78:1311–1323.

Fryxell, J. M., C. Packer, K. McCann, E. J. Solberg, and B.-E. Sæther. 2010. Resource management cycles and the sustainability of harvested wildlife populations. Science 328:903–906.

Fulton, E. A., A. D. Smith, D. C. Smith, and I. E. van Putten. 2011. Human behaviour: the key source of uncertainty in fisheries management. Fish and Fisheries 12:2–17.

Gamelon, M., B. K. Sandercock, and B. E. Sæther. 2019. Does harvesting amplify environmentally induced population fluctuations over time in marine and terrestrial species? Journal of Applied Ecology 56:2186–2194.

Goto, D., A. A. Filin, D. Howell, B. Bogstad, Y. Kovalev, and H. Gjøsæter. 2021. Tradeoffs of managing cod as a sustainable resource in fluctuating environments. Ecological Applications xx:xx–xx.

Harwood, J., and K. Stokes. 2003. Coping with uncertainty in ecological advice: lessons from fisheries. Trends in Ecology & Evolution 18:617–622.

Hilborn, R., R. O. Amoroso, C. M. Anderson, J. K. Baum, T. A. Branch, C. Costello, C. L. de Moor, A. Faraj, D. Hively, and O. P. Jensen. 2020. Effective fisheries management instrumental in improving fish stock status. Proceedings of the National Academy of Sciences.

Hilborn, R., J.-J. Maguire, A. M. Parma, and A. A. Rosenberg. 2001. The precautionary approach and risk management: can they increase the probability of successes in fishery management? Canadian Journal of Fisheries 58:99–107.

Hordyk, A. R., Q. C. Huynh, and T. R. Carruthers. 2019. Misspecification in stock assessments: Common uncertainties and asymmetric risks. Fish and Fisheries 20:888–902.

Hurtado-Ferro, F., C. S. Szuwalski, J. L. Valero, S. C. Anderson, C. J. Cunningham, K. F. Johnson, R. Licandeo, C. R. McGilliard, C. C. Monnahan, and M. L. Muradian. 2015. Looking in the rear-view mirror: bias and retrospective patterns in integrated, age-structured stock assessment models. ICES Journal of Marine Science 72:99–110.

ICES. 2015. Report of the Joint ICES-MYFISH Workshop to consider the basis for FMSY ranges for all stocks (WKMSYREF3), 17–21 November 2014, Charlottenlund, Denmark. ICES CM 2014/ACOM:64. 156 pp.

ICES. 2016. Report of the Benchmark Workshop on North Sea Stocks (WKNSEA), 14–18 March 2016, Copenhagen, Denmark. ICES CM 2016/ACOM:37 704 pp.

ICES. 2017. Report of the Workshop to consider FMSY ranges for stocks in ICES categories 1 and 2 in Western Waters (WKMSYREF4), 13–16 October 2015, Brest, France.

ICES. 2018. Report of the Working Group on the Assessment of Demersal Stocks in the North Sea and Skagerrak (WGNSSK), 24 April-3 May 2018, Oostende, Belgium. ICES CM 2018/ACOM:22. 1264 pp.

ICES. 2019a. Report of the Interbenchmark Protocol on North Sea Saithe. (IBPNSsaithe). ICES Scientific Reports. VOL 1:ISS 1. 65 pp.

ICES. 2019b. Workshop on guidelines for management strategy evaluations (WKGMSE2). ICES Scientific Reports. 1:33. 162 pp.

ICES. 2019c. Workshop on north sea stocks management strategy evaluation (WKNSMSE). ICES Scientific Reports. 1:12. 378 pp.

ICES. 2020a. The third Workshop on Guidelines for Management Strategy Evaluations (WKGMSE3). ICES Scientific Reports. 2:116. 112 pp.

ICES. 2020b. Workshop on Catch Forecast from Biased Assessments (WKFORBIAS; outputs from 2019 meeting). ICES Scientific Reports. 2:28. 38 pp.

ICES. 2020c. Workshop on Management Strategy Evaluation of Mackerel (WKMSEMAC). ICES Scientific Reports. 2:74. 175 pp.

ICES. 2021. ICES fisheries management reference points for category 1 and 2 stocks. ICES Technical Guidelines.

Kell, L., M. Pastoors, R. Scott, M. Smith, F. Van Beek, C. O’Brien, and G. Pilling. 2005. Evaluation of multiple management objectives for Northeast Atlantic flatfish stocks: sustainability vs. stability of yield. ICES Journal of Marine Science 62:1104–1117.

Kell, L., G. Pilling, G. Kirkwood, M. Pastoors, B. Mesnil, K. Korsbrekke, P. Abaunza, R. Aps, A. Biseau, and P. Kunzlik. 2006. An evaluation of multi-annual management strategies for ICES roundfish stocks. ICES Journal of Marine Science 63:12–24.

Kell, L. T., I. Mosqueira, P. Grosjean, J.-M. Fromentin, D. Garcia, R. Hillary, E. Jardim, S. Mardle, M. Pastoors, and J. Poos. 2007. FLR: an open-source framework for the evaluation and development of management strategies. ICES Journal of Marine Science 64:640–646.

Kraak, S., F. Buisman, M. Dickey-Collas, J. Poos, M. Pastoors, J. Smit, J. Van Oostenbrugge, and N. Daan. 2008. The effect of management choices on the sustainability and economic performance of a mixed fishery: a simulation study. ICES Journal of Marine Science 65:697–712.

Kuparinen, A., D. M. Keith, and J. A. Hutchings. 2014. Allee effect and the uncertainty of population recovery. Conservation Biology 28:790–798.

Mangel, M., L. M. Talbot, G. K. Meffe, M. T. Agardy, D. L. Alverson, J. Barlow, D. B. Botkin, G. Budowski, T. Clark, and J. Cooke. 1996. Principles for the conservation of wild living resources. Ecological Applications 6:338–362.

Memarzadeh, M., and C. Boettiger. 2018. Adaptive management of ecological systems under partial observability. Biological Conservation 224:9–15.

Memarzadeh, M., G. L. Britten, B. Worm, and C. Boettiger. 2019. Rebuilding global fisheries under uncertainty. Proceedings of the National Academy of Sciences 116:15985–15990.

Milner-Gulland, E., K. Shea, H. Possingham, T. Coulson, and C. Wilcox. 2001. Competing harvesting strategies in a simulated population under uncertainty. Animal Conservation 4:157–167.

Nielsen, A., and C. W. Berg. 2014. Estimation of time-varying selectivity in stock assessments using state-space models. Fisheries Research 158:96–101.

Ovaskainen, O., and B. Meerson. 2010. Stochastic models of population extinction. Trends in Ecology & Evolution 25:643–652.

Patterson, K., R. Cook, C. Darby, S. Gavaris, L. Kell, P. Lewy, B. Mesnil, A. Punt, V. Restrepo, and D. W. Skagen. 2001. Estimating uncertainty in fish stock assessment and forecasting. Fish and Fisheries 2:125–157.

Patterson, K., and M. Résimont. 2007. Change and stability in landings: the responses of fisheries to scientific advice and TACs. ICES Journal of Marine Science 64:714–717.

Punt, A. E., D. S. Butterworth, C. L. de Moor, J. A. A. De Oliveira, and M. Haddon. 2016. Management strategy evaluation: best practices. Fish and Fisheries 17:303–334.

Punt, A. E., G. N. Tuck, J. Day, C. M. Canales, J. M. Cope, C. L. de Moor, J. A. De Oliveira, M. Dickey-Collas, B. Þ. Elvarsson, and M. A. Haltuch. 2020. When are model-based stock assessments rejected for use in management and what happens then? Fisheries Research 224:105465.

Rindorf, A., M. Cardinale, S. Shephard, J. A. De Oliveira, E. Hjorleifsson, A. Kempf, A. Luzenczyk, C. Millar, D. C. Miller, and C. L. Needle10. 2016. Fishing for MSY: can “pretty good yield” ranges be used without impairing recruitment. ICES Journal of Marine Science.

Ripa, J., and P. Lundberg. 1996. Noise colour and the risk of population extinctions. Proceedings of the Royal Society of London B: Biological Sciences 263:1751–1753.

Sethi, S. A. 2010. Risk management for fisheries. Fish and Fisheries 11:341–365.

Smith, A., K. Sainsbury, and R. Stevens. 1999. Implementing effective fisheries-management systems–management strategy evaluation and the Australian partnership approach. ICES Journal of Marine Science 56:967–979.

Szuwalski, C. S., J. N. Ianelli, and A. E. Punt. 2017. Reducing retrospective patterns in stock assessment and impacts on management performance. ICES Journal of Marine Science 75:596–609.

UN. 2015. Transforming our world: the 2030 Agenda for Sustainable Development A/RES/70/1. Division for Sustainable Development Goals: New York, NY, USA.

Walters, C., and J.-J. Maguire. 1996. Lessons for stock assessment from the northern cod collapse. Reviews in Fish Biology and Fisheries 6:125–137.

Walters, C. J., and R. Hilborn. 1978. Ecological optimization and adaptive management. Annual review of Ecology and Systematics 9:157–188.

Wiedenmann, J., and O. P. Jensen. 2018. Uncertainty in stock assessment estimates for New England groundfish and its impact on achieving target harvest rates. Canadian Journal of Fisheries and Aquatic Sciences 75:342–356.

Wiedenmann, J., M. J. Wilberg, A. Sylvia, and T. J. Miller. 2015. Autocorrelated error in stock assessment estimates: implications for management strategy evaluation. Fisheries Research 172:325–334.

