## Appendix S1 for "Shaping sustainable harvest boundaries for marine populations despite estimation bias"

**Appendix S1**. Input data and parameter estimates for the North Sea saithe popualtion operating model used in management strategy evaluation

Table S1. Age-specific landings numbers (thousands) of North Sea saithe in 1967–2017.

| year | age 3 | age 4 | age 5 | age 6 | age 7 | age 8 | age 9 | age 10 and older |
| --- | --- | --- | --- | --- | --- | --- | --- | --- |
| 1967 | 17330 | 16220 | 15531 | 2303 | 1594 | 292 | 198 | 183 |
| 1968 | 23223 | 21231 | 13184 | 6023 | 429 | 242 | 123 | 145 |
| 1969 | 30235 | 17681 | 11057 | 7609 | 5738 | 791 | 626 | 150 |
| 1970 | 37249 | 76661 | 15000 | 12128 | 3894 | 1792 | 318 | 267 |
| 1971 | 69808 | 57792 | 32737 | 4736 | 4248 | 2843 | 1874 | 774 |
| 1972 | 48075 | 66095 | 25317 | 21207 | 3672 | 2944 | 1641 | 1607 |
| 1973 | 54332 | 37698 | 26849 | 16061 | 8428 | 2000 | 1357 | 2381 |
| 1974 | 66938 | 33740 | 14123 | 20688 | 14666 | 5199 | 1477 | 1955 |
| 1975 | 56987 | 25864 | 10319 | 7566 | 13657 | 9357 | 3501 | 2687 |
| 1976 | 207823 | 53060 | 11696 | 6253 | 3976 | 5362 | 3586 | 3490 |
| 1977 | 27461 | 54967 | 14755 | 5490 | 3777 | 3447 | 3812 | 4701 |
| 1978 | 35059 | 27269 | 18062 | 3312 | 1138 | 1033 | 768 | 3484 |
| 1979 | 16332 | 14216 | 11182 | 8699 | 2805 | 733 | 540 | 2089 |
| 1980 | 17494 | 12341 | 9015 | 6718 | 5658 | 1150 | 509 | 2302 |
| 1981 | 26178 | 8339 | 6739 | 3675 | 3335 | 3396 | 657 | 2536 |
| 1982 | 31895 | 40587 | 9174 | 5978 | 2145 | 1454 | 982 | 1254 |
| 1983 | 28242 | 20604 | 26013 | 5678 | 4893 | 1494 | 1036 | 1327 |
| 1984 | 80933 | 32172 | 12957 | 13011 | 1657 | 1252 | 335 | 646 |
| 1985 | 134024 | 55605 | 13281 | 4765 | 3005 | 682 | 399 | 742 |
| 1986 | 55435 | 91223 | 15186 | 5381 | 2603 | 1456 | 445 | 900 |
| 1987 | 31220 | 97470 | 13990 | 3158 | 1811 | 1240 | 910 | 700 |
| 1988 | 32578 | 26408 | 35323 | 3828 | 1908 | 1104 | 776 | 680 |
| 1989 | 22128 | 30752 | 13187 | 10951 | 1557 | 739 | 419 | 488 |
| 1990 | 40808 | 19583 | 11322 | 4714 | 2776 | 745 | 281 | 364 |
| 1991 | 46117 | 29871 | 7467 | 3583 | 1716 | 953 | 367 | 458 |
| 1992 | 18404 | 33614 | 12753 | 3193 | 1524 | 696 | 518 | 422 |
| 1993 | 37823 | 20828 | 11845 | 3125 | 1568 | 1511 | 814 | 1026 |
| 1994 | 19958 | 40193 | 13034 | 4297 | 947 | 346 | 427 | 794 |
| 1995 | 26664 | 26034 | 14797 | 3774 | 3494 | 674 | 552 | 800 |
| 1996 | 11066 | 38861 | 11786 | 7731 | 3163 | 808 | 210 | 491 |
| 1997 | 15036 | 19299 | 30177 | 3676 | 2640 | 1012 | 291 | 288 |
| 1998 | 10363 | 31017 | 16367 | 16077 | 2231 | 1206 | 567 | 277 |
| 1999 | 9429 | 13872 | 26684 | 8389 | 10070 | 2346 | 891 | 657 |
| 2000 | 7064 | 17295 | 8940 | 12339 | 3159 | 3226 | 641 | 441 |
| 2001 | 16052 | 17646 | 22421 | 3349 | 3586 | 1772 | 1614 | 245 |
| 2002 | 9131 | 31779 | 12286 | 13307 | 2245 | 2220 | 1199 | 1479 |
| 2003 | 13009 | 24646 | 20397 | 6836 | 6855 | 1535 | 2000 | 2042 |
| 2004 | 8037 | 20071 | 15649 | 15220 | 2037 | 2164 | 1300 | 1066 |
| 2005 | 9191 | 15473 | 19060 | 20042 | 7956 | 1628 | 1188 | 1151 |
| 2006 | 12200 | 26690 | 9986 | 11286 | 8395 | 3824 | 1008 | 1281 |
| 2007 | 15181 | 10163 | 19157 | 7078 | 5564 | 3610 | 1218 | 930 |
| 2008 | 6924 | 23230 | 10930 | 14196 | 4977 | 3276 | 3551 | 3118 |
| 2009 | 6607 | 14349 | 13827 | 5817 | 8419 | 2978 | 1505 | 2934 |
| 2010 | 7880 | 8859 | 9174 | 6394 | 2670 | 4762 | 1679 | 2669 |
| 2011 | 10150 | 22799 | 8852 | 3630 | 2860 | 1183 | 1563 | 3869 |
| 2012 | 7029 | 11712 | 15572 | 4016 | 1971 | 1267 | 537 | 2610 |
| 2013 | 4999 | 25516 | 4974 | 7645 | 1886 | 1241 | 616 | 1658 |
| 2014 | 3099 | 12117 | 13380 | 3737 | 4047 | 1036 | 429 | 1388 |
| 2015 | 6206 | 7392 | 13555 | 8021 | 1844 | 1621 | 715 | 975 |
| 2016 | 3508 | 10374 | 8756 | 5156 | 2732 | 1423 | 852 | 1317 |
| 2017 | 3033 | 15139 | 8795 | 6179 | 5362 | 1876 | 820 | 1111 |

Table S2. Age-specific discards numbers (thousands) of North Sea saithe in 1967–2017.

| year | age 3 | age 4 | age 5 | age 6 | age 7 | age 8 | age 9 | age 10 and older |
| --- | --- | --- | --- | --- | --- | --- | --- | --- |
| 1967 | 9617 | 3175 | 1141 | 55 | 16 | 7 | 5 | 2 |
| 1968 | 12888 | 4156 | 969 | 143 | 4 | 6 | 3 | 2 |
| 1969 | 16779 | 3461 | 813 | 181 | 57 | 19 | 16 | 2 |
| 1970 | 20671 | 15007 | 1102 | 288 | 38 | 42 | 8 | 3 |
| 1971 | 38741 | 11313 | 2406 | 112 | 42 | 67 | 48 | 9 |
| 1972 | 26680 | 12938 | 1861 | 504 | 36 | 69 | 42 | 18 |
| 1973 | 30152 | 7380 | 1973 | 381 | 83 | 47 | 35 | 26 |
| 1974 | 37148 | 6605 | 1038 | 491 | 144 | 122 | 38 | 22 |
| 1975 | 31626 | 5063 | 758 | 180 | 135 | 220 | 89 | 30 |
| 1976 | 115333 | 10387 | 860 | 148 | 39 | 126 | 92 | 38 |
| 1977 | 15240 | 10760 | 1084 | 130 | 37 | 81 | 97 | 52 |
| 1978 | 19456 | 5338 | 1327 | 79 | 11 | 24 | 20 | 38 |
| 1979 | 9063 | 2783 | 822 | 207 | 28 | 17 | 14 | 23 |
| 1980 | 9709 | 2416 | 662 | 160 | 56 | 27 | 13 | 25 |
| 1981 | 14527 | 1632 | 495 | 87 | 33 | 80 | 17 | 28 |
| 1982 | 17700 | 7945 | 674 | 142 | 21 | 34 | 25 | 14 |
| 1983 | 15673 | 4033 | 1912 | 135 | 48 | 35 | 26 | 15 |
| 1984 | 44915 | 6298 | 952 | 309 | 16 | 29 | 9 | 7 |
| 1985 | 74378 | 10885 | 976 | 113 | 30 | 16 | 10 | 8 |
| 1986 | 30764 | 17857 | 1116 | 128 | 26 | 34 | 11 | 10 |
| 1987 | 17326 | 19080 | 1028 | 75 | 18 | 29 | 23 | 8 |
| 1988 | 18079 | 5169 | 2596 | 91 | 19 | 26 | 20 | 7 |
| 1989 | 12280 | 6020 | 969 | 260 | 15 | 17 | 11 | 5 |
| 1990 | 22647 | 3833 | 832 | 112 | 27 | 18 | 7 | 4 |
| 1991 | 25593 | 5847 | 549 | 85 | 17 | 22 | 9 | 5 |
| 1992 | 10213 | 6580 | 937 | 76 | 15 | 16 | 13 | 5 |
| 1993 | 20990 | 4077 | 871 | 74 | 15 | 36 | 21 | 11 |
| 1994 | 11076 | 7868 | 958 | 102 | 9 | 8 | 11 | 9 |
| 1995 | 14797 | 5096 | 1087 | 90 | 34 | 16 | 14 | 9 |
| 1996 | 6141 | 7607 | 866 | 184 | 31 | 19 | 5 | 5 |
| 1997 | 8344 | 3778 | 2218 | 87 | 26 | 24 | 7 | 3 |
| 1998 | 5751 | 6072 | 1203 | 382 | 22 | 28 | 14 | 3 |
| 1999 | 5233 | 2716 | 1961 | 199 | 99 | 55 | 23 | 7 |
| 2000 | 3920 | 3386 | 657 | 293 | 31 | 76 | 16 | 5 |
| 2001 | 8908 | 3454 | 1648 | 80 | 35 | 42 | 41 | 3 |
| 2002 | 8439 | 5710 | 2451 | 425 | 64 | 324 | 121 | 96 |
| 2003 | 15288 | 7106 | 234 | 0 | 0 | 0 | 0 | 0 |
| 2004 | 5605 | 4407 | 0 | 0 | 0 | 0 | 0 | 0 |
| 2005 | 3498 | 0 | 0 | 0 | 0 | 0 | 0 | 0 |
| 2006 | 5114 | 5282 | 394 | 0 | 0 | 0 | 0 | 0 |
| 2007 | 9433 | 3152 | 1762 | 97 | 0 | 0 | 0 | 0 |
| 2008 | 696 | 7682 | 1610 | 745 | 111 | 9 | 0 | 0 |
| 2009 | 831 | 1158 | 395 | 30 | 93 | 16 | 14 | 11 |
| 2010 | 886 | 390 | 266 | 117 | 1 | 11 | 0 | 38 |
| 2011 | 2636 | 1470 | 129 | 44 | 7 | 25 | 1 | 8 |
| 2012 | 7305 | 1341 | 1377 | 58 | 7 | 1 | 4 | 1 |
| 2013 | 2268 | 4801 | 339 | 224 | 4 | 0 | 0 | 1 |
| 2014 | 955 | 2205 | 1816 | 220 | 77 | 4 | 0 | 1 |
| 2015 | 2163 | 931 | 704 | 232 | 17 | 3 | 0 | 2 |
| 2016 | 3874 | 3867 | 905 | 573 | 26 | 7 | 1 | 0 |
| 2017 | 1943 | 3850 | 978 | 69 | 2 | 0 | 0 | 2 |

Table S3. Age-specific mass (kg) of North Sea saithe in 1967–2017.

| year | age 3 | | age 4 | | age 5 | | age 6 | | age 7 | | age 8 | | age 9 | | age 10 and older |
| --- | --- | --- | --- | --- | --- | --- | --- | --- | --- | --- | --- | --- | --- | --- | --- |
| 1967 | 0.898 | 1.339 | | 2.094 | | 3.183 | | 3.753 | | 5.316 | | 5.891 | | 7.719 | |
| 1968 | 1.234 | 1.624 | | 1.979 | | 3.007 | | 4.039 | | 4.428 | | 6.136 | | 7.406 | |
| 1969 | 0.933 | 1.530 | | 2.251 | | 2.711 | | 3.558 | | 4.406 | | 5.220 | | 6.767 | |
| 1970 | 0.908 | 1.416 | | 2.049 | | 2.716 | | 3.599 | | 4.463 | | 5.687 | | 6.845 | |
| 1971 | 0.811 | 1.325 | | 2.167 | | 2.934 | | 3.765 | | 4.634 | | 5.172 | | 6.163 | |
| 1972 | 0.780 | 1.175 | | 1.952 | | 2.367 | | 3.793 | | 4.228 | | 4.630 | | 6.326 | |
| 1973 | 0.792 | 1.382 | | 1.633 | | 2.569 | | 3.356 | | 4.684 | | 4.814 | | 6.445 | |
| 1974 | 0.831 | 1.534 | | 2.372 | | 2.751 | | 3.428 | | 4.498 | | 5.713 | | 7.857 | |
| 1975 | 0.862 | 1.472 | | 2.479 | | 3.298 | | 3.764 | | 4.296 | | 5.540 | | 7.562 | |
| 1976 | 0.678 | 1.287 | | 2.250 | | 3.068 | | 4.034 | | 4.383 | | 5.112 | | 7.147 | |
| 1977 | 0.733 | 1.234 | | 1.926 | | 3.108 | | 4.161 | | 4.605 | | 4.859 | | 6.542 | |
| 1978 | 0.793 | 1.304 | | 2.145 | | 3.338 | | 4.521 | | 4.900 | | 5.449 | | 7.400 | |
| 1979 | 1.069 | 1.595 | | 2.228 | | 3.093 | | 4.049 | | 5.274 | | 6.308 | | 7.955 | |
| 1980 | 0.921 | 1.790 | | 2.380 | | 3.028 | | 4.089 | | 5.126 | | 5.939 | | 8.148 | |
| 1981 | 0.927 | 1.790 | | 2.705 | | 3.584 | | 4.535 | | 5.478 | | 6.980 | | 8.724 | |
| 1982 | 1.048 | 1.548 | | 2.518 | | 3.218 | | 4.206 | | 5.125 | | 5.905 | | 8.823 | |
| 1983 | 0.992 | 1.688 | | 2.139 | | 3.135 | | 3.690 | | 4.632 | | 5.505 | | 8.453 | |
| 1984 | 0.767 | 1.586 | | 2.286 | | 2.688 | | 3.895 | | 4.665 | | 6.183 | | 8.474 | |
| 1985 | 0.640 | 1.244 | | 1.941 | | 2.769 | | 3.406 | | 4.950 | | 5.865 | | 8.854 | |
| 1986 | 0.670 | 1.018 | | 1.786 | | 2.430 | | 3.571 | | 4.209 | | 5.651 | | 8.218 | |
| 1987 | 0.650 | 0.861 | | 1.815 | | 3.072 | | 4.209 | | 5.330 | | 6.128 | | 8.603 | |
| 1988 | 0.752 | 0.964 | | 1.379 | | 2.789 | | 4.023 | | 5.254 | | 6.322 | | 8.649 | |
| 1989 | 0.864 | 1.018 | | 1.413 | | 1.997 | | 3.913 | | 5.017 | | 6.430 | | 8.431 | |
| 1990 | 0.815 | 1.175 | | 1.575 | | 2.245 | | 3.241 | | 4.858 | | 6.315 | | 8.416 | |
| 1991 | 0.764 | 1.138 | | 1.744 | | 2.363 | | 3.165 | | 4.222 | | 6.066 | | 8.191 | |
| 1992 | 0.930 | 1.169 | | 1.599 | | 2.240 | | 3.667 | | 4.330 | | 5.412 | | 7.045 | |
| 1993 | 0.868 | 1.239 | | 1.746 | | 2.634 | | 3.184 | | 3.980 | | 5.080 | | 6.891 | |
| 1994 | 0.911 | 1.100 | | 1.594 | | 2.432 | | 3.617 | | 4.787 | | 6.548 | | 8.326 | |
| 1995 | 0.967 | 1.272 | | 1.807 | | 2.560 | | 3.554 | | 4.767 | | 5.267 | | 7.891 | |
| 1996 | 0.933 | 1.167 | | 1.798 | | 2.366 | | 2.951 | | 4.705 | | 6.092 | | 8.382 | |
| 1997 | 0.873 | 1.125 | | 1.445 | | 2.585 | | 3.555 | | 4.525 | | 6.158 | | 8.866 | |
| 1998 | 0.861 | 0.949 | | 1.386 | | 1.743 | | 2.948 | | 3.883 | | 4.996 | | 7.227 | |
| 1999 | 0.850 | 1.042 | | 1.206 | | 1.752 | | 2.337 | | 3.493 | | 4.844 | | 6.745 | |
| 2000 | 0.992 | 1.107 | | 1.532 | | 1.683 | | 2.593 | | 3.084 | | 4.773 | | 7.461 | |
| 2001 | 0.774 | 1.053 | | 1.307 | | 2.093 | | 2.546 | | 3.485 | | 4.141 | | 6.141 | |
| 2002 | 0.776 | 1.014 | | 1.495 | | 1.791 | | 2.961 | | 3.761 | | 4.638 | | 5.750 | |
| 2003 | 0.636 | 0.889 | | 1.167 | | 1.810 | | 2.368 | | 3.176 | | 3.768 | | 5.065 | |
| 2004 | 0.794 | 1.010 | | 1.392 | | 1.896 | | 2.860 | | 3.687 | | 4.814 | | 7.059 | |
| 2005 | 0.715 | 1.155 | | 1.325 | | 1.710 | | 2.132 | | 3.026 | | 3.622 | | 5.713 | |
| 2006 | 0.904 | 1.012 | | 1.489 | | 1.906 | | 2.424 | | 3.058 | | 4.318 | | 5.734 | |
| 2007 | 0.769 | 1.124 | | 1.286 | | 1.834 | | 2.328 | | 2.887 | | 3.600 | | 4.975 | |
| 2008 | 0.916 | 1.065 | | 1.488 | | 1.692 | | 2.210 | | 2.792 | | 3.206 | | 4.565 | |
| 2009 | 1.033 | 1.333 | | 1.672 | | 1.994 | | 2.566 | | 3.086 | | 3.651 | | 4.790 | |
| 2010 | 1.037 | 1.474 | | 2.033 | | 2.597 | | 3.163 | | 3.488 | | 3.968 | | 5.223 | |
| 2011 | 0.955 | 1.192 | | 1.787 | | 2.571 | | 3.068 | | 3.418 | | 3.718 | | 4.289 | |
| 2012 | 0.910 | 1.287 | | 1.383 | | 2.196 | | 3.221 | | 3.536 | | 4.181 | | 4.482 | |
| 2013 | 0.878 | 1.132 | | 1.586 | | 1.957 | | 3.076 | | 3.841 | | 4.541 | | 5.648 | |
| 2014 | 1.091 | 1.265 | | 1.568 | | 2.334 | | 2.607 | | 4.010 | | 5.530 | | 6.679 | |
| 2015 | 0.951 | 1.253 | | 1.621 | | 2.180 | | 3.037 | | 3.793 | | 4.228 | | 7.285 | |
| 2016 | 0.937 | 1.239 | | 1.611 | | 2.231 | | 2.888 | | 3.450 | | 4.331 | | 6.208 | |
| 2017 | 0.956 | 1.228 | | 1.755 | | 2.356 | | 2.987 | | 4.232 | | 4.473 | | 6.287 | |

| year | age 3 | age 4 | age 5 | age 6 | age 7 | age 8 | age 9 | age 10 and older |
| --- | --- | --- | --- | --- | --- | --- | --- | --- |
| 1967 | 0.931 | 1.362 | 2.104 | 3.186 | 3.754 | 5.316 | 5.891 | 7.719 |
| 1968 | 1.278 | 1.652 | 1.989 | 3.009 | 4.040 | 4.428 | 6.136 | 7.406 |
| 1969 | 0.966 | 1.557 | 2.261 | 2.713 | 3.559 | 4.406 | 5.220 | 6.768 |
| 1970 | 0.941 | 1.441 | 2.059 | 2.718 | 3.600 | 4.463 | 5.687 | 6.845 |
| 1971 | 0.840 | 1.348 | 2.178 | 2.936 | 3.766 | 4.634 | 5.173 | 6.163 |
| 1972 | 0.808 | 1.196 | 1.961 | 2.369 | 3.794 | 4.228 | 4.630 | 6.326 |
| 1973 | 0.821 | 1.406 | 1.641 | 2.571 | 3.357 | 4.684 | 4.814 | 6.445 |
| 1974 | 0.861 | 1.561 | 2.383 | 2.753 | 3.429 | 4.498 | 5.713 | 7.857 |
| 1975 | 0.893 | 1.498 | 2.490 | 3.300 | 3.765 | 4.296 | 5.540 | 7.562 |
| 1976 | 0.702 | 1.309 | 2.260 | 3.071 | 4.035 | 4.383 | 5.112 | 7.147 |
| 1977 | 0.760 | 1.256 | 1.935 | 3.111 | 4.162 | 4.605 | 4.859 | 6.542 |
| 1978 | 0.822 | 1.327 | 2.155 | 3.340 | 4.522 | 4.901 | 5.449 | 7.400 |
| 1979 | 1.107 | 1.623 | 2.238 | 3.095 | 4.050 | 5.274 | 6.308 | 7.955 |
| 1980 | 0.955 | 1.821 | 2.391 | 3.030 | 4.090 | 5.126 | 5.939 | 8.148 |
| 1981 | 0.961 | 1.821 | 2.718 | 3.587 | 4.536 | 5.478 | 6.980 | 8.724 |
| 1982 | 1.086 | 1.575 | 2.529 | 3.220 | 4.207 | 5.125 | 5.905 | 8.823 |
| 1983 | 1.028 | 1.718 | 2.149 | 3.138 | 3.691 | 4.632 | 5.505 | 8.453 |
| 1984 | 0.795 | 1.614 | 2.297 | 2.690 | 3.896 | 4.665 | 6.183 | 8.474 |
| 1985 | 0.663 | 1.265 | 1.951 | 2.772 | 3.407 | 4.950 | 5.865 | 8.854 |
| 1986 | 0.694 | 1.035 | 1.794 | 2.432 | 3.572 | 4.209 | 5.651 | 8.218 |
| 1987 | 0.674 | 0.876 | 1.824 | 3.075 | 4.210 | 5.330 | 6.128 | 8.603 |
| 1988 | 0.779 | 0.981 | 1.386 | 2.791 | 4.024 | 5.254 | 6.322 | 8.649 |
| 1989 | 0.895 | 1.036 | 1.420 | 1.998 | 3.914 | 5.018 | 6.430 | 8.431 |
| 1990 | 0.844 | 1.196 | 1.583 | 2.247 | 3.242 | 4.858 | 6.315 | 8.416 |
| 1991 | 0.791 | 1.158 | 1.752 | 2.365 | 3.165 | 4.222 | 6.066 | 8.191 |
| 1992 | 0.964 | 1.189 | 1.607 | 2.242 | 3.668 | 4.330 | 5.413 | 7.046 |
| 1993 | 0.899 | 1.260 | 1.754 | 2.636 | 3.185 | 3.980 | 5.080 | 6.891 |
| 1994 | 0.944 | 1.119 | 1.601 | 2.434 | 3.618 | 4.787 | 6.548 | 8.326 |
| 1995 | 1.002 | 1.294 | 1.816 | 2.562 | 3.555 | 4.767 | 5.267 | 7.891 |
| 1996 | 0.967 | 1.187 | 1.807 | 2.368 | 2.952 | 4.705 | 6.092 | 8.382 |
| 1997 | 0.905 | 1.145 | 1.452 | 2.587 | 3.556 | 4.525 | 6.158 | 8.866 |
| 1998 | 0.892 | 0.966 | 1.393 | 1.744 | 2.949 | 3.883 | 4.996 | 7.227 |
| 1999 | 0.881 | 1.061 | 1.211 | 1.754 | 2.337 | 3.493 | 4.844 | 6.745 |
| 2000 | 1.027 | 1.127 | 1.539 | 1.684 | 2.594 | 3.084 | 4.773 | 7.462 |
| 2001 | 0.802 | 1.072 | 1.313 | 2.095 | 2.546 | 3.485 | 4.141 | 6.141 |
| 2002 | 0.923 | 1.035 | 1.478 | 1.769 | 2.947 | 3.426 | 4.407 | 5.674 |
| 2003 | 0.833 | 0.980 | 1.173 | 1.810 | 2.368 | 3.176 | 3.768 | 5.065 |
| 2004 | 0.918 | 1.084 | 1.392 | 1.896 | 2.860 | 3.687 | 4.814 | 7.059 |
| 2005 | 0.921 | 1.155 | 1.325 | 1.710 | 2.132 | 3.026 | 3.622 | 5.713 |
| 2006 | 0.945 | 1.069 | 1.514 | 1.906 | 2.424 | 3.058 | 4.318 | 5.734 |
| 2007 | 0.837 | 1.143 | 1.317 | 1.840 | 2.328 | 2.887 | 3.600 | 4.975 |
| 2008 | 0.944 | 1.193 | 1.565 | 1.720 | 2.226 | 2.795 | 3.206 | 4.565 |
| 2009 | 1.036 | 1.340 | 1.664 | 1.992 | 2.563 | 3.085 | 3.648 | 4.793 |
| 2010 | 1.036 | 1.479 | 2.034 | 2.597 | 3.164 | 3.488 | 3.968 | 5.199 |
| 2011 | 1.007 | 1.207 | 1.783 | 2.573 | 3.068 | 3.404 | 3.717 | 4.284 |
| 2012 | 1.015 | 1.321 | 1.408 | 2.201 | 3.223 | 3.536 | 4.177 | 4.482 |
| 2013 | 0.898 | 1.156 | 1.614 | 1.976 | 3.078 | 3.841 | 4.541 | 5.648 |
| 2014 | 1.126 | 1.300 | 1.607 | 2.384 | 2.617 | 4.013 | 5.530 | 6.679 |
| 2015 | 0.977 | 1.244 | 1.625 | 2.190 | 3.043 | 3.796 | 4.228 | 7.287 |
| 2016 | 0.998 | 1.292 | 1.628 | 2.283 | 2.892 | 3.453 | 4.333 | 6.208 |
| 2017 | 1.047 | 1.302 | 1.809 | 2.361 | 2.988 | 4.232 | 4.473 | 6.292 |

Table S4. Age-specific mass (kg) of North Sea saithe landings in 1967–2017.

Table S5. Age-specific maturity (proportion of aduts) of North Sea saithe.

|  | age 1 | age 2 | age 3 | age 4 | age 5 | age 6 | age 7 | age 8 and older |
| --- | --- | --- | --- | --- | --- | --- | --- | --- |
| Maturity | 0.00 | 0.00 | 0.00 | 0.20 | 0.65 | 0.84 | 0.97 | 1.00 |

| year | CPUE |
| --- | --- |
| 2000 | 2.104 |
| 2001 | 2.356 |
| 2002 | 1.926 |
| 2003 | 1.794 |
| 2004 | 2.275 |
| 2005 | 2.491 |
| 2006 | 2.584 |
| 2007 | 2.148 |
| 2008 | 2.583 |
| 2009 | 2.026 |
| 2010 | 1.888 |
| 2011 | 1.871 |
| 2012 | 1.641 |
| 2013 | 1.800 |
| 2014 | 1.751 |
| 2015 | 1.973 |
| 2016 | 1.725 |
| 2017 | 1.970 |

Table S6. Aggregate exploitable biomass index (catch per unit effort or CPUE) of North Sea saithe in 2000–2017.

Table S7. Age-specific abundunce indices of North Sea saithe in 1992–2017.

| year | age 3 | age 4 | age 5 | age 6 | age 7 | age 8 |
| --- | --- | --- | --- | --- | --- | --- |
| 1992 | 1.077 | 2.760 | 0.516 | 0.098 | 0.057 | 0.050 |
| 1993 | 7.965 | 2.781 | 1.129 | 0.197 | 0.011 | 0.040 |
| 1994 | 1.117 | 1.615 | 0.893 | 0.609 | 0.091 | 0.040 |
| 1995 | 13.959 | 2.501 | 1.559 | 0.533 | 0.172 | 0.049 |
| 1996 | 3.825 | 6.533 | 1.112 | 0.971 | 0.212 | 0.069 |
| 1997 | 3.756 | 3.351 | 7.461 | 0.698 | 0.534 | 0.181 |
| 1998 | 1.181 | 4.134 | 1.351 | 1.580 | 0.149 | 0.179 |
| 1999 | 2.086 | 1.907 | 3.155 | 0.619 | 0.632 | 0.074 |
| 2000 | 3.479 | 8.836 | 1.081 | 0.868 | 0.114 | 0.152 |
| 2001 | 21.475 | 6.169 | 3.936 | 0.356 | 0.444 | 0.113 |
| 2002 | 10.748 | 18.974 | 1.327 | 1.090 | 0.162 | 0.264 |
| 2003 | 19.272 | 23.802 | 13.402 | 0.393 | 0.439 | 0.168 |
| 2004 | 4.930 | 6.727 | 3.237 | 0.921 | 0.064 | 0.085 |
| 2005 | 8.916 | 7.512 | 4.428 | 1.914 | 1.082 | 0.104 |
| 2006 | 10.553 | 29.579 | 2.835 | 1.177 | 0.445 | 0.242 |
| 2007 | 34.006 | 5.578 | 11.700 | 1.016 | 0.743 | 0.358 |
| 2008 | 3.312 | 5.584 | 0.907 | 1.997 | 0.254 | 0.254 |
| 2009 | 1.346 | 1.703 | 0.568 | 0.101 | 0.229 | 0.200 |
| 2010 | 1.361 | 0.964 | 0.471 | 0.205 | 0.045 | 0.166 |
| 2011 | 4.520 | 8.451 | 1.059 | 1.114 | 0.426 | 0.080 |
| 2012 | 11.134 | 2.497 | 2.968 | 0.503 | 0.483 | 0.344 |
| 2013 | 14.701 | 16.279 | 1.830 | 1.858 | 0.308 | 0.146 |
| 2014 | 1.649 | 3.923 | 2.822 | 0.481 | 0.520 | 0.114 |
| 2015 | 11.001 | 5.613 | 4.611 | 1.581 | 0.289 | 0.285 |
| 2016 | 37.901 | 17.439 | 3.255 | 2.681 | 0.945 | 0.195 |
| 2017 | 11.447 | 13.102 | 3.068 | 1.267 | 0.942 | 0.473 |
