## Appendix S2 for "Shaping sustainable harvest boundaries for marine populations despite estimation bias"

**Appendix S2**. Details of model fitting for the North Sea saithe popuation operating model used in management strategy evaluation and additional analyses using alterantive model specification.

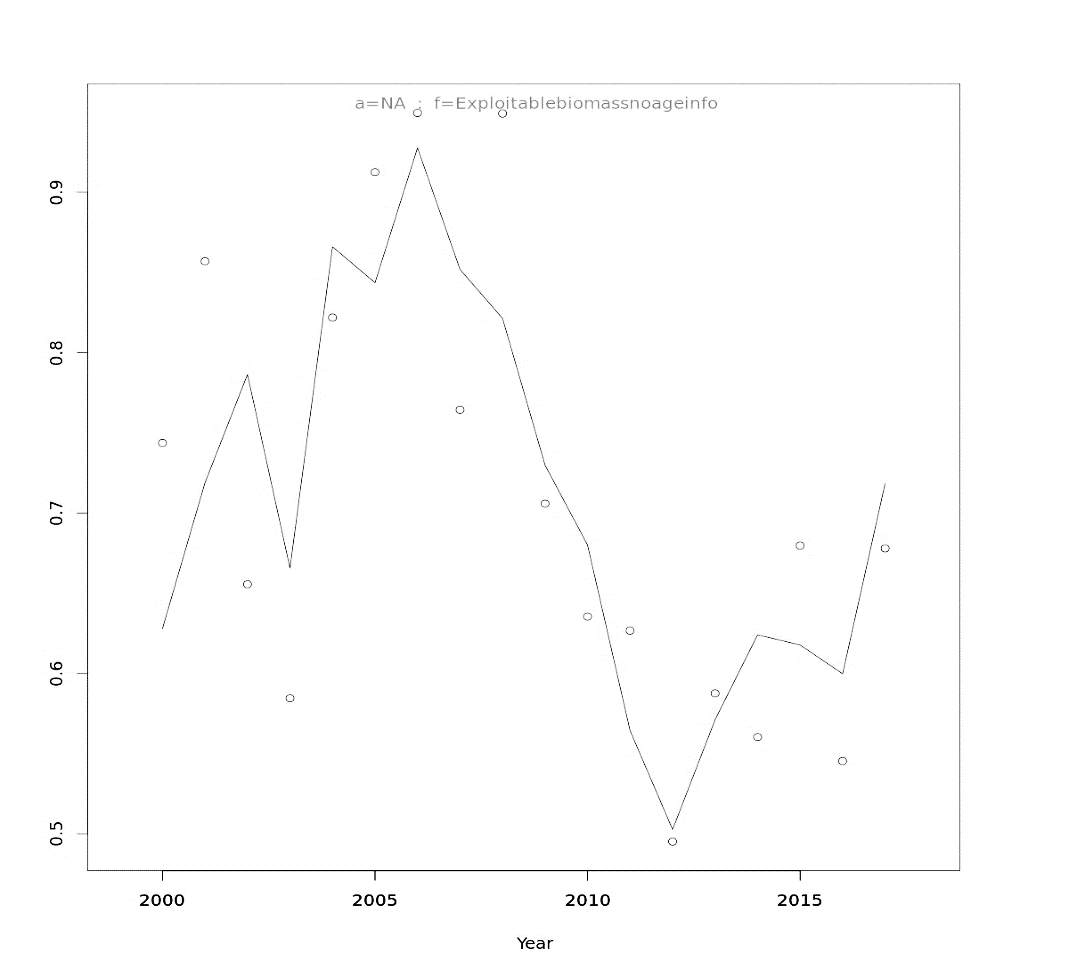
Figure S1. Fit to the time series of commercial catch data (age-aggregated biomass of German, French, and Norwegian trawlers) in 2000–2017 using State-space Assessment Model.

Figure S2. Fit to the time series of age-specific (three to eight) abundance indices (International bottom trawl surveys in the third quarter, IBTS-Q3) in 1992–2017 using State-space Assessment Model.

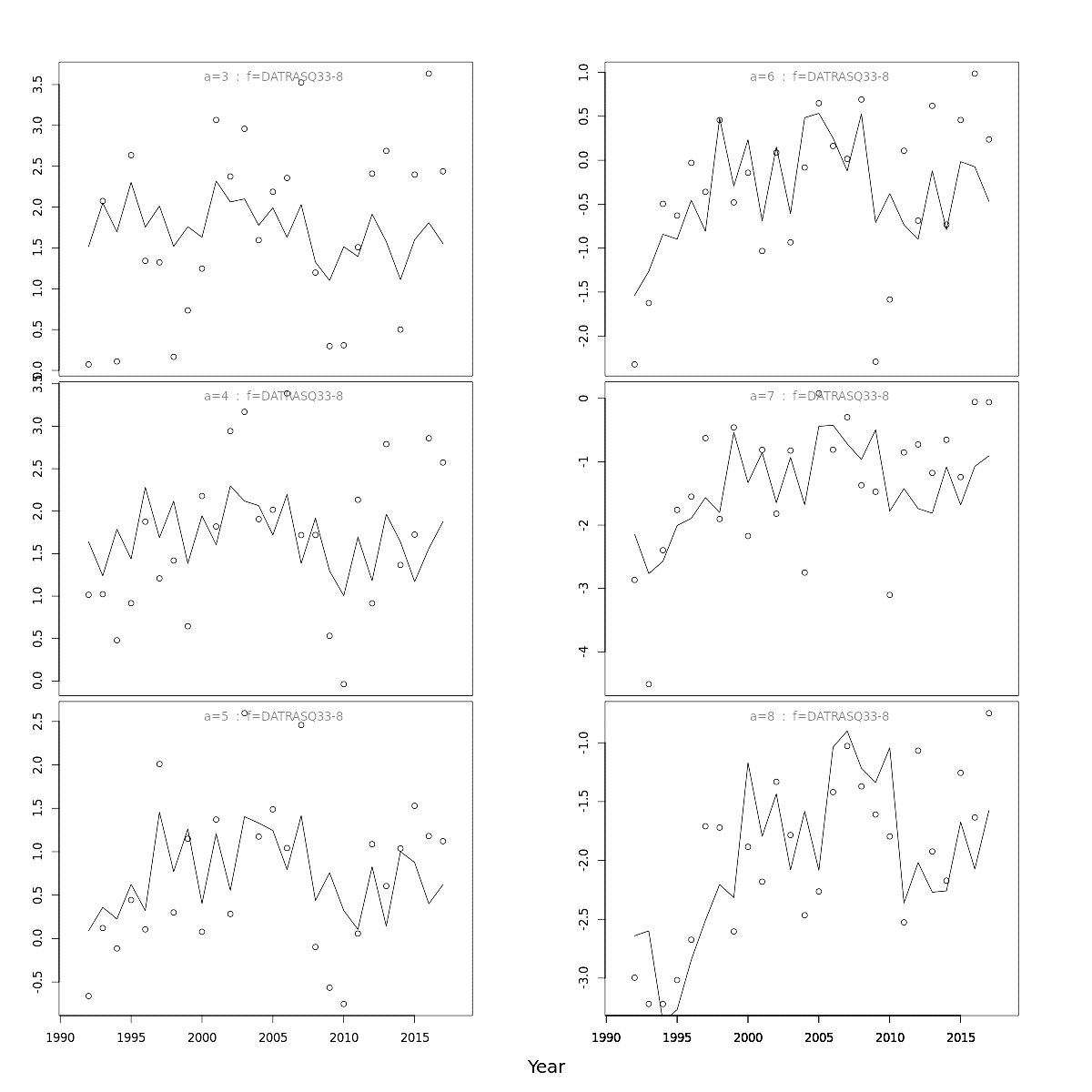

Age 3

Age 4

Age 5

Age 6

Age 7

Age 8

Figure S3. Process and sampling errors in the North Sea saithe population and observation operating models. (a) recruitment and survival process error and (b) fishing mortality process error estimated from normalized residuals from a state-space estimation model in the 2018 assessment. Standardized residuals for total catch (c), monitoring survey (IBTS–Q3, d), and exploitable biomass index (e) from the 2018 assessment. Blue circles indicate a positive residual
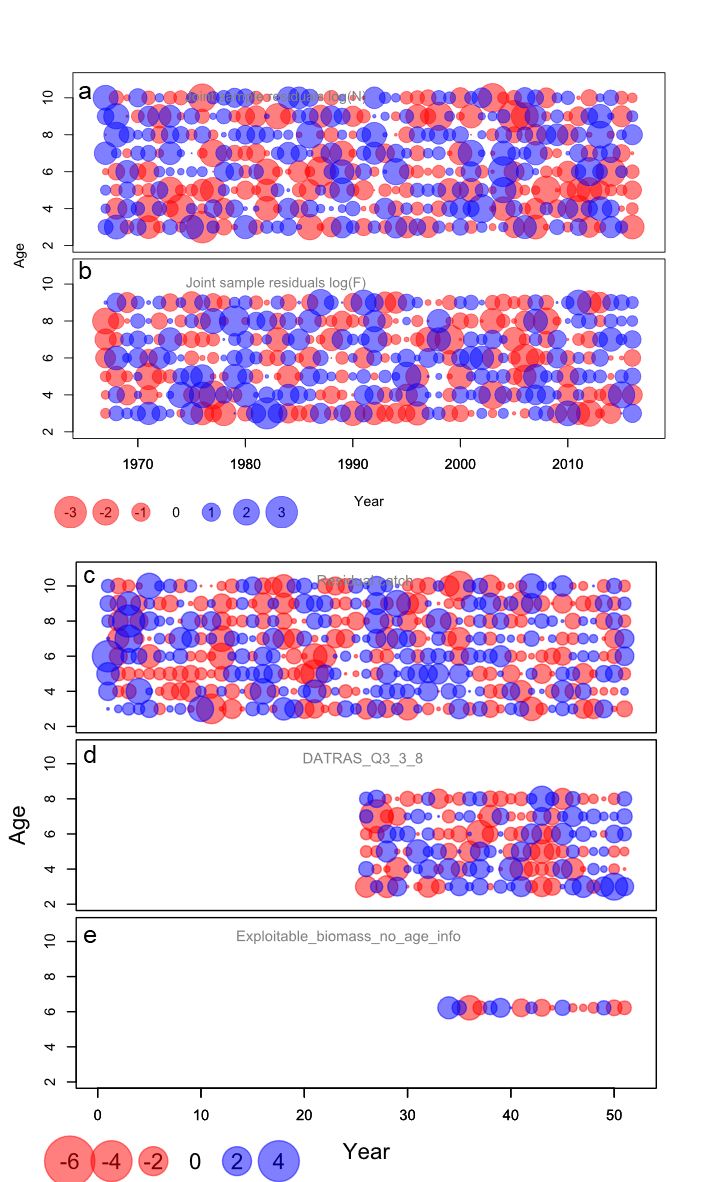
and red circles a negative residual.

Figure S4. Recruitment in the North Sea saithe operating model. (a) spawner-recruit pairs from historical data (red circles) and fitted values (black circles) generated from the stock-recruit model (dark gray solid lines) for randomly selected 100 replicates. (b) historical (red line) and simulated recruits (black line) using empirical cumulative distribution function (ecdf in R) for the stock-recruit pairs in a. (c) autocorrelation in the 1998–2017 recruit estimates from the 2018 assessment.

Figure S5. Monitoring and catch surveys in the North Sea saithe operating model. (a) observed (IBTS–Q3) and simulated age-specific number survey indices in 1992–2017. (b) observed and simulated aggregated exploitable biomass index in 2000
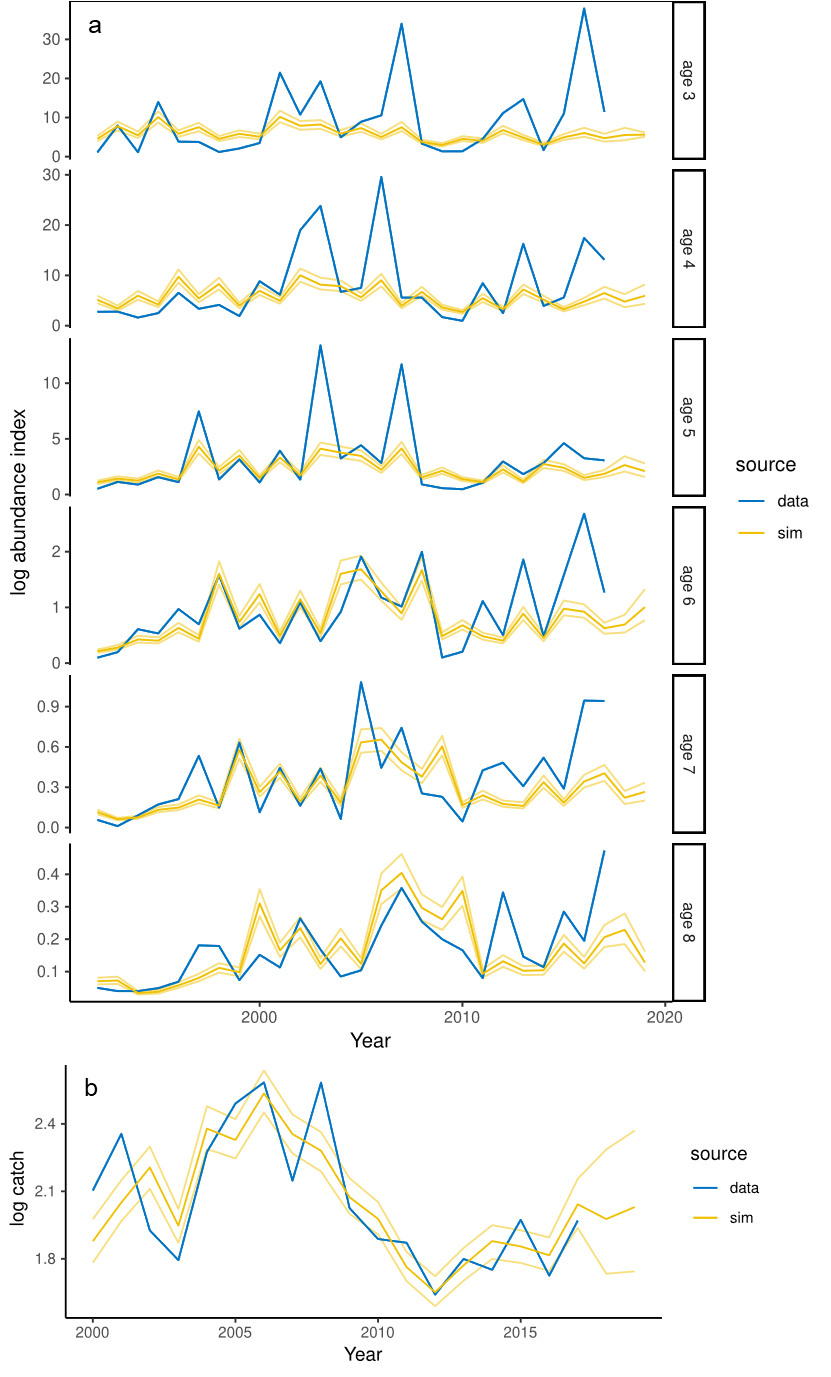
–2017. For simulated indices, a thick line indicates median, thin lines indicate 75% confidence limits.

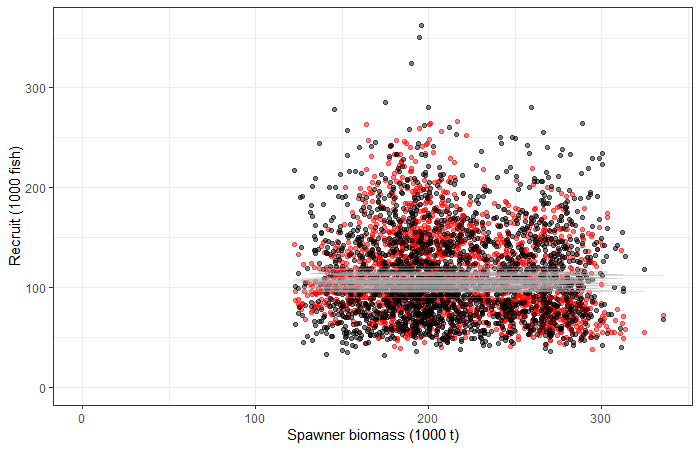

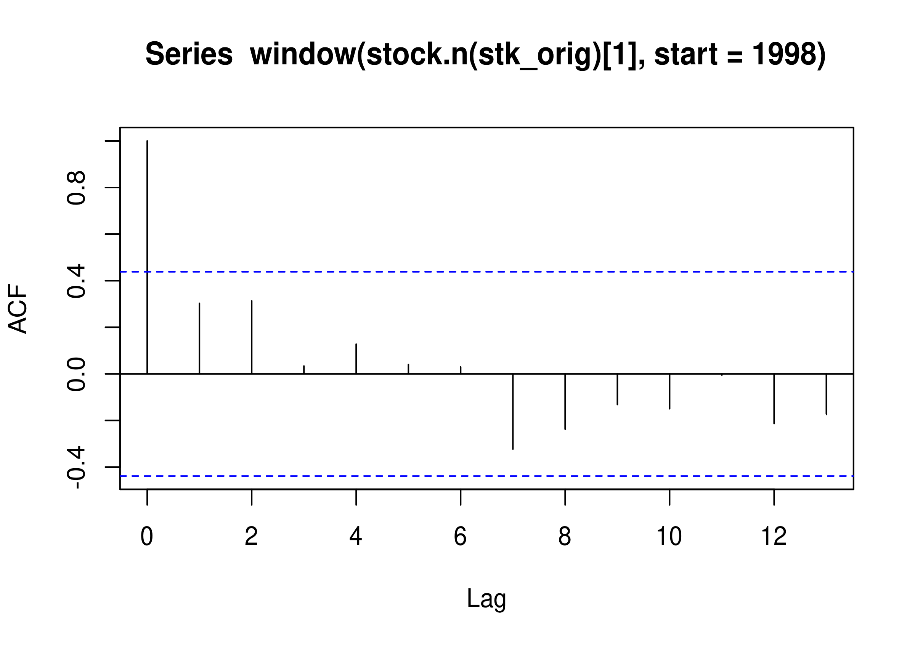

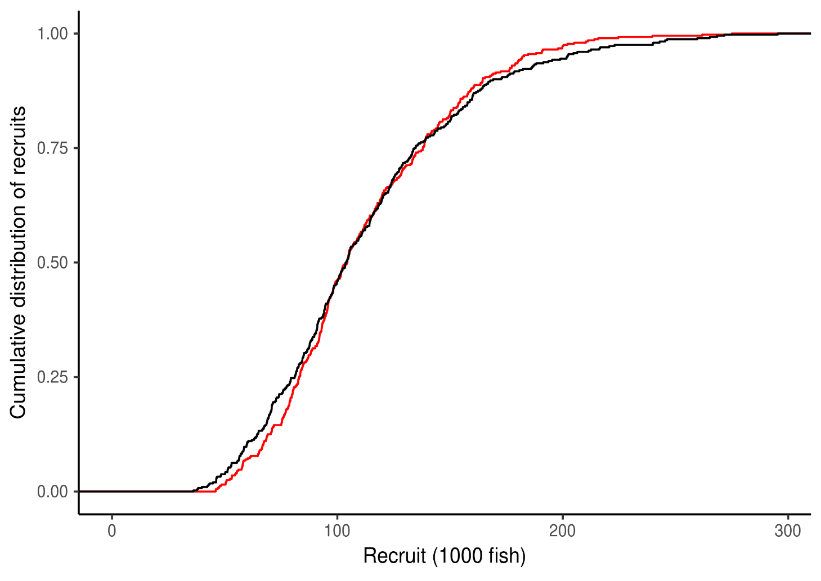

c

a

b

Figure S6. Estimated correlations between age groups for (a) monitoring survey (IBTS Q3) index-at-age and (b) catch-at-age (no correlations modelled).

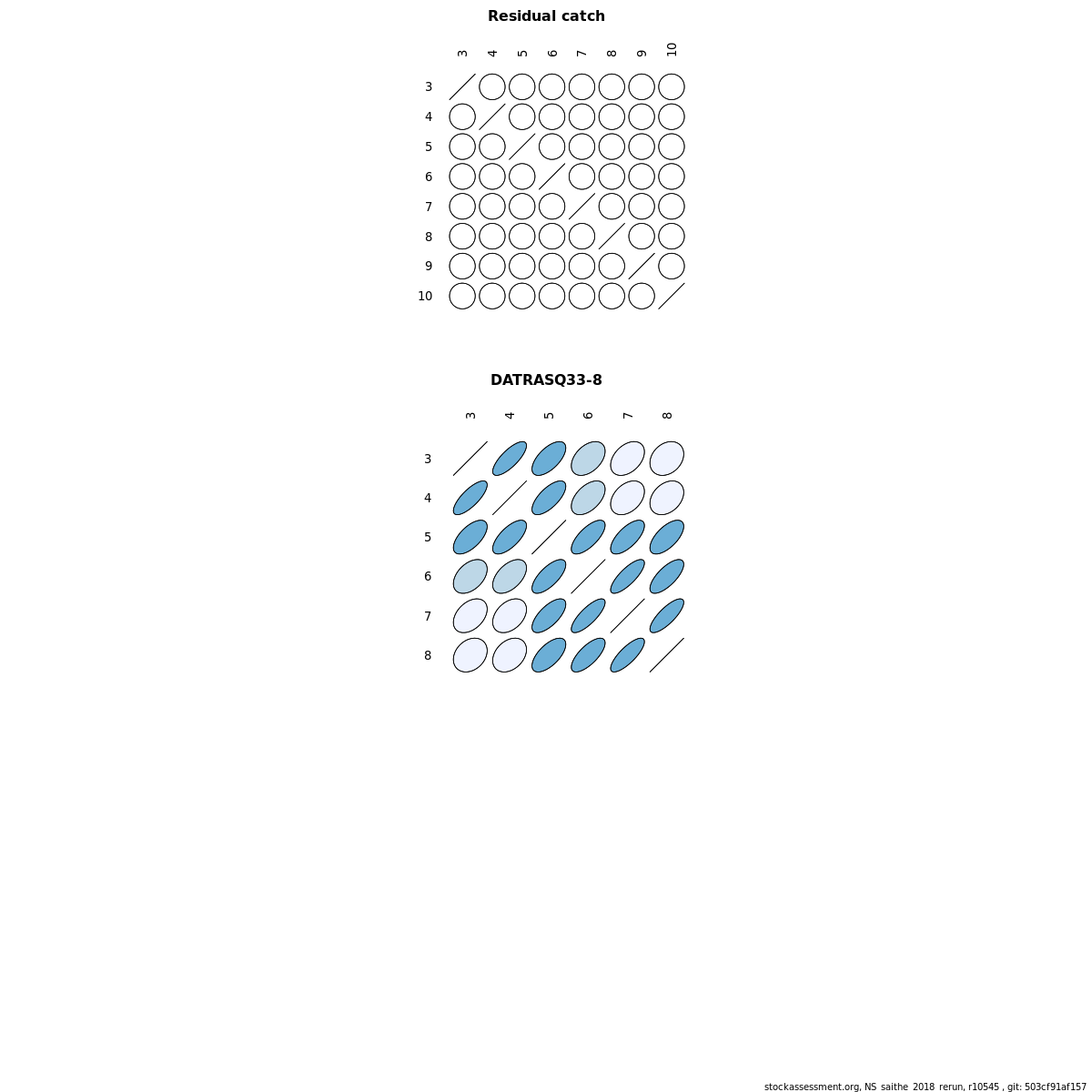

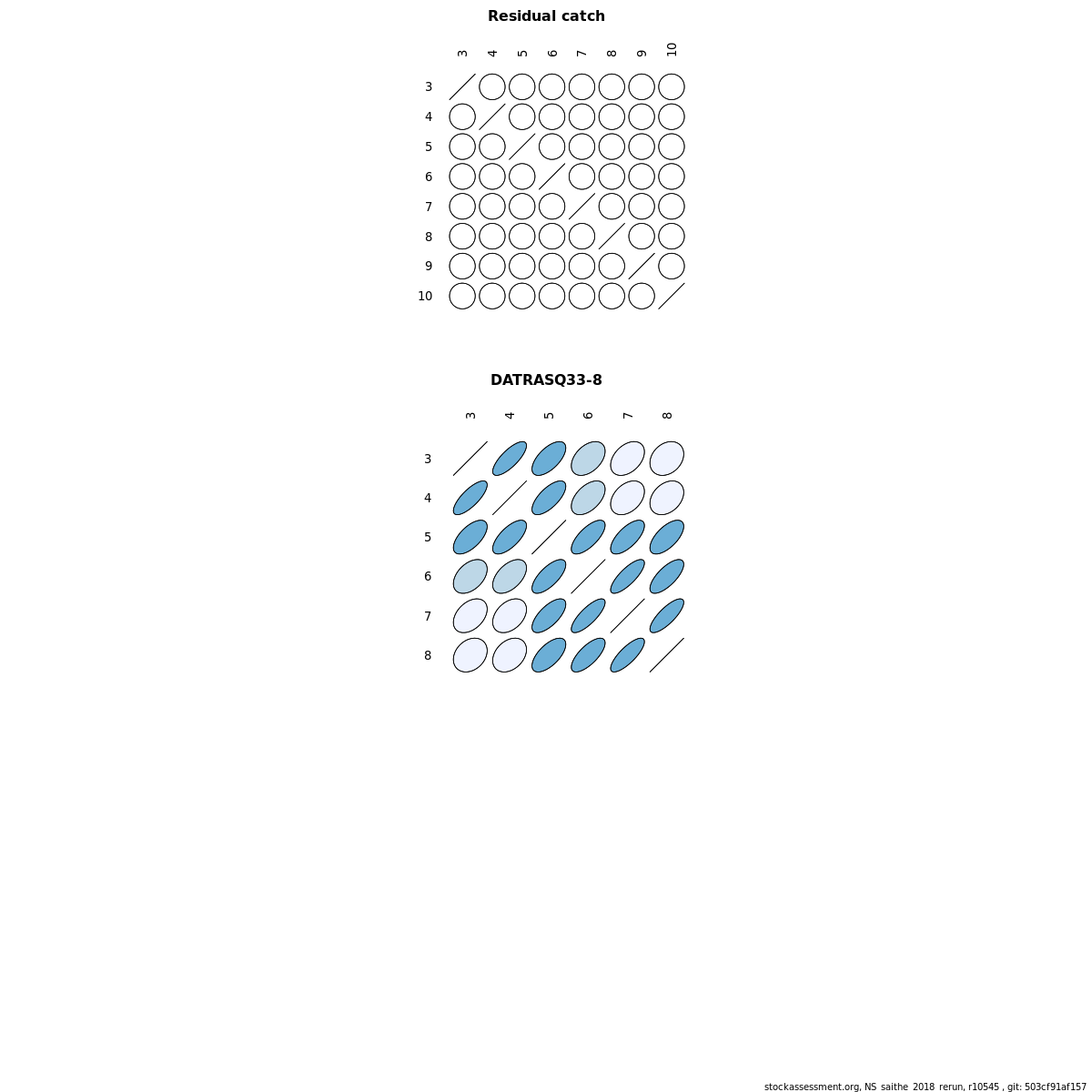

a

b

Figure S7. Projected population and harvest dynamics of North Sea saithe under the baseline (no bias) scenario with the initial control parameter values prior to optimization in the 2019 management strategy evaluation (*F*_target_ = *F*_MSY_ = 0.363 and *B*_trigger_ = MSY *B*_trigger_ = 149,098 t). Top plot is recruitment (three-year-olds), second plot SSB, third plot catch, and bottom plot mean *F* (four to seven-year-olds). The vertical black line separates the historical period from the projection period. The SSB plot includes *B*_pa_ = MSY *B_t_*_rigger_ (horizontal solid line) and *B*_lim_ (horizontal hashed line), while the mean *F* plot includes *F*_MSY_ (horizontal solid line) and *F*_lim_ (limit reference point for fishing mortality rate computed with Eqsim, horizontal dashed line). The plots show medians (solid black line) with the darker shaded area indicating the 25th and 75th percentiles, and the light shaded area the 5th and 95th percentiles. The coloured lines represent the values from five replicates (replicates 100-105).

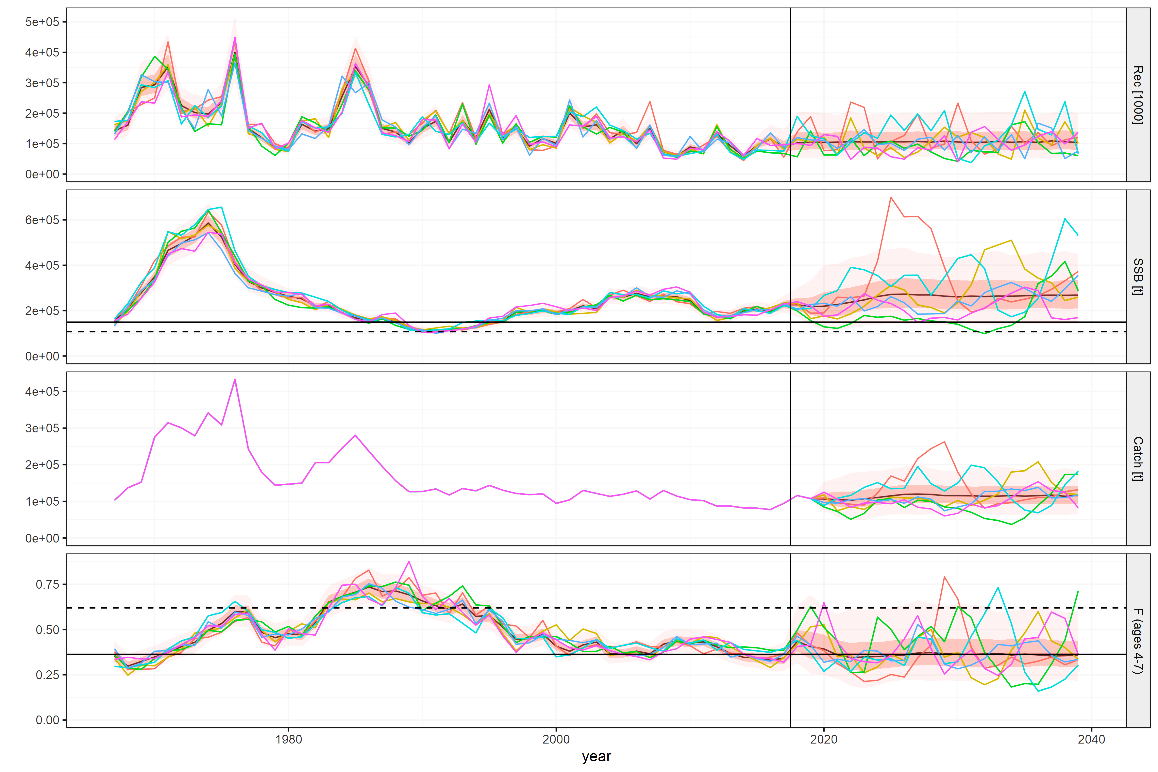

Figure S8. Projected population and harvest dynamics of North Sea saithe under the baseline (no bias) scenario with the optimized control parameter values in the 2019 management strategy evaluation (*F*_target_ = 0.35 and *B*_trigger_ = 250,000 t). Top plot is recruitment (three-year-olds), second plot SSB, third plot catch, and bottom plot mean *F* (four to seven-year-olds). The vertical black line separates the historical period from the projection period. The SSB plot includes *B*_pa_ = MSY *B_t_*_rigger_ (horizontal solid line) and *B*_lim_ (horizontal hashed line), while the mean *F* plot includes *F*_MSY_ (horizontal solid line) and *F*_lim_ (limit reference point for fishing mortality rate computed with Eqsim, horizontal dashed line). The plots show medians (solid black line) with the darker shaded area indicating the 25th and 75th percentiles, and the light shaded area the 5th and 95th percentiles. The coloured lines represent the values from five replicates (replicates 100-105).

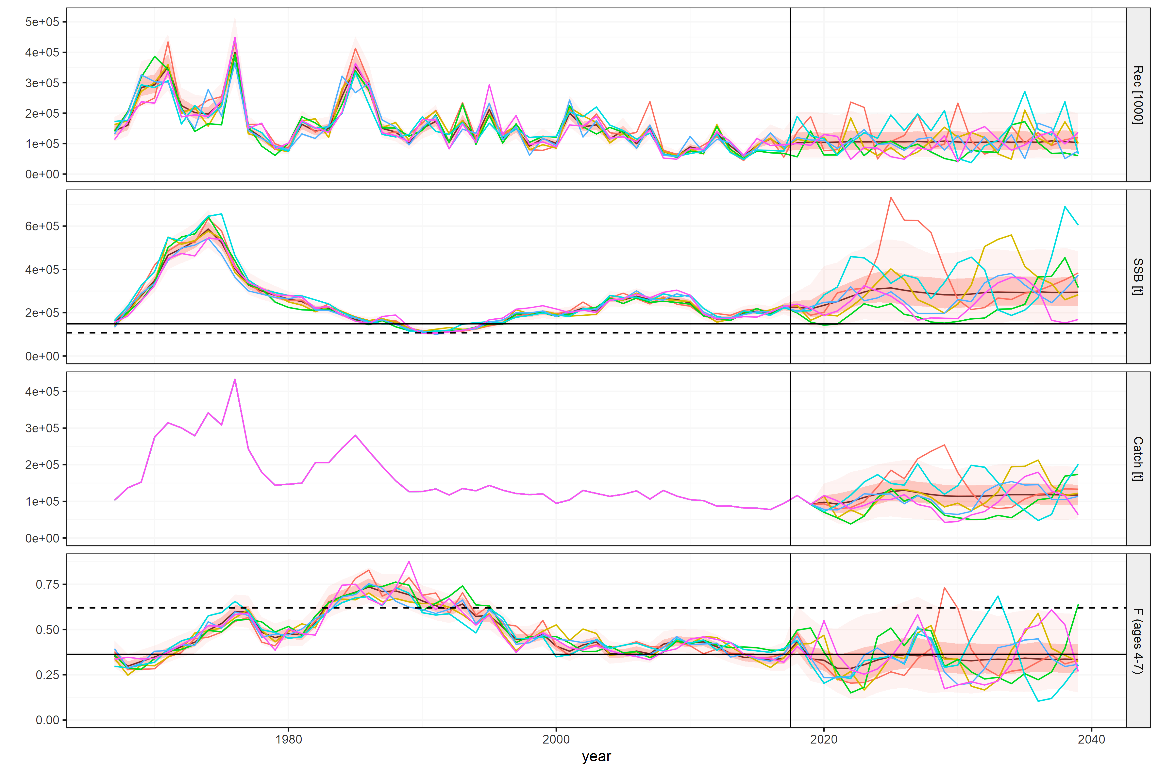

Figure S9. Estimates of fishing pressure (*F*) and adult biomass (SSB) from the management procedure (MP) compared to the underlying “truth” for alternative operating models (OMs) with natural mortality rates, *M* = 0.2 (baseline, top panel), *M* = 0.1 (overestimated in the MP, middle), and *M* = 0.3 (underestimated in the MP, bottom). Values > 1 indicate an overestimation by the management procedure while values < 1 indicate an underestimation. The plots show medians (solid black line) with the darker shaded area indicating the 25th and 75th percentiles, and the light shaded area the 5th and 95th percentiles.

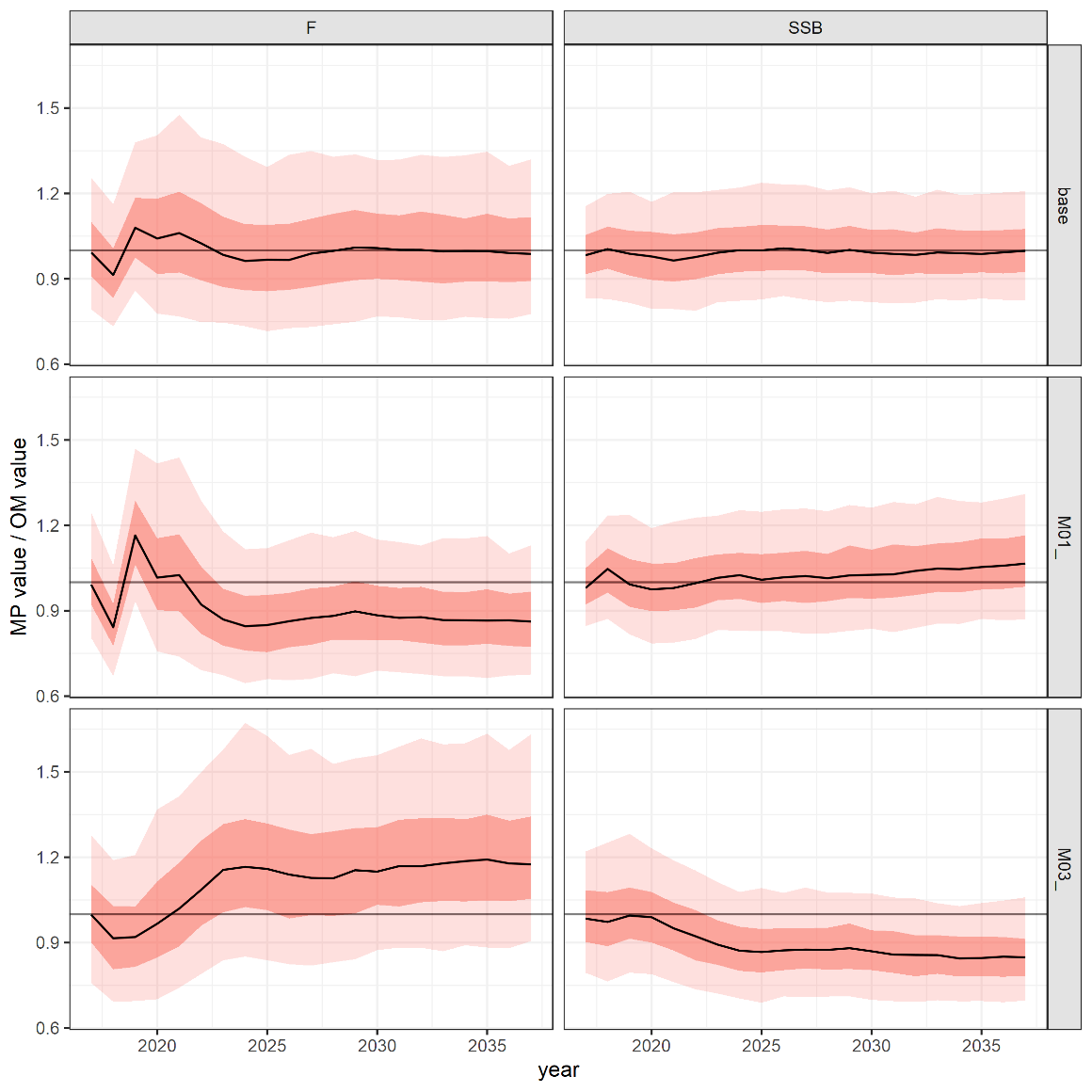
